# Silc1 long noncoding RNA is an immediate-early gene promoting efficient memory formation

**DOI:** 10.1101/2022.11.11.516100

**Authors:** Rotem Ben-Tov Perry, Michael Tsoory, Michael Tolmasov, Igor Ulitsky

## Abstract

Long noncoding RNAs (lncRNAs) are expressed in many brain circuits and neuronal types, but their significance to normal brain functions has remained largely unknown. Here, we study the functions in the central nervous system of *Silc1*, a lncRNA we previously showed to be important for neuroregeneration in the peripheral nervous system. We found that *Silc1* is rapidly and strongly induced upon stimulation in the hippocampus and is required for efficient spatial learning. *Silc1* production is important for the induction of *Sox11* (its cis-regulated target gene) throughout the CA1-CA3 regions and the proper expression of key *Sox11* target genes. Consistent with its newly found role in neuronal plasticity, we find that during aging and in models of Alzheimer’s disease *Silc1* levels decline. Overall, we uncover a novel plasticity pathway, in which *Silc1* acts as an immediate-early gene to activate *Sox11* to induce a neuronal growth-associated transcriptional program important for memory formation.

## Introduction

Long noncoding RNAs (lncRNAs) are products of pervasive transcription of eukaryotic genomes. While tens of thousands of lncRNA genes have now been annotated in the mammalian genomes (Frankish et al. 2021), the functions of the vast majority of them, if any, remain unclear. Cells of the nervous system express a particularly rich repertoire of lncRNA genes, and some were indicated to play particularly important roles in neurogenesis and/or the functioning of the nervous system (reviewed in (Hezroni, Perry, and Ulitsky 2019)). Some of those are also affected by various neurological diseases.

The molecular mechanisms underlying memory formation and retrieval remain only partially understood. Some lncRNAs have been implicated in the process via analysis of genetic models. Loss of the *Carip* lncRNA, which binds specifically to CaMKIIβ, affects the phosphorylation of AMPA and NMDA receptors and causes dysfunction of synaptic transmission and attenuated long-term potentiation, which results in impaired spatial memory formation (Cui et al. 2022). Other lncRNAs, such as Neat1, were studied using transient perturbations and shown to affect memory formation (Butler et al. 2019)(Hezroni, Perry, and Ulitsky 2019).

We have recently characterized lncRNA expression during regeneration in the peripheral nervous system and characterized two lncRNAs induced by sciatic nerve crush and regulating neurite outgrowth (Perry et al. 2018). We have further shown that one of these, *Silc1*, a lncRNA conserved in sequence throughout mammals, is required for timely regeneration *in vivo*, and its loss in *Silc1^-/-^* mice was associated with reduced expression of *Sox11*, a transcription factor with well-established roles in neurogenesis and in neuroregeneration, both in the adult brain and in the regenerating dorsal root ganglia (DRG) (Tsang, Oliemuller, and Howard 2020; Perry et al. 2018). *Silc1* is transcribed from within a large gene desert flanking the *Sox11* gene, which lies ~200 kb upstream of it, and the two loci appear in spatial proximity to each other in various chromatin capture datasets (Perry et al. 2018; Malysheva et al. 2018; Bonev et al. 2017). The human ortholog, *SILC1*, was recently studied in neuroblastoma cells, where SOX11 is a lineage-dependence factor (Decaesteker et al. 2020; Ye et al. 2019).

*Sox11*, a member of the SoxC family transcription factors, has so far been primarily studied in the context of embryonic neurogenesis, where it was proposed to have overlapping targets with other SoxC transcription factors (TFs) *Sox4* and *Sox12*, as well as with other members of the Sox family (Bergsland et al. 2011). *Sox11^-/-^* mice die shortly after birth (Sock et al. 2004), consistent with its requirement for both proliferation and growth of neuronal cells of various types (Lin et al. 2011; Potzner et al. 2010; Thein et al. 2010). Conditional loss of *Sox11* in the neuronal lineage (obtained using tamoxifen-inducible Nestin-driven Cre) demonstrated that loss of *Sox11* reduces neurogenesis in both the embryo and the adult (Y. Wang et al. 2013). Specific ablation of *Sox11* in the subgranular zone (SGZ) of the dentate gyrus (DG) in the hippocampus reduced the number of DCX+ or NeuroD1+ cells (Y. Wang et al. 2013). In addition to proliferation and differentiation, loss of Sox11 also affects axonal growth in embryonic sensory neurons *in vivo* and *in vitro* (Lin et al. 2011) and axonal growth in the adult DRG neurons upon injury (Jankowski, Miller, and Koerber 2018).

Little is known about the roles of *Sox11* in the post-mitotic neurons in the brain, where although its expression is modest, it is one of the most abundantly expressed Sox genes. *Sox11* levels in the adult brain are particularly high in the SGZ of the adult hippocampus (Haslinger et al. 2009; Mu et al. 2012), which is one of the two sites of ongoing adult neurogenesis (Ming and Song 2005). At early postnatal stages, *Sox11* is particularly high in the late neuroblasts, and immature granule and pyramidal neurons, and then its levels decline in mature neurons (Hochgerner et al. 2018). *Sox11* mRNA is broadly induced in the DG upon electroconvulsive stimulation (Sun et al. 2005; Su et al. 2017), suggesting a role in neuronal activity. A recent study has further studied this increase and noted that it also occurs when mice are placed in a novel environment, specifically in mature neurons in the granule layer of the DG (von Wittgenstein et al. 2020), where its expression is sparse while mice are housed under standard conditions. This “novelty”-induced increase was associated with a subset of the Fos-positive cells (von Wittgenstein et al. 2020) which experienced stronger neuronal activation. Notably, SOX11-positive cells were not observed in the CA subfields in that study. Several target genes of SOX11 are known, including *Dcx*, which is expressed in a tightly overlapping domain with *Sox11* (Mu et al. 2012).

As it is presently impossible to deduce the function of lncRNAs from their sequences or structures, co-expression with annotated protein-coding genes is often used as a first and readily available method to predict the function of long noncoding RNAs (Guttman et al. 2011, 2009; Yan et al. 2015). However, it remains unclear whether lncRNAs are necessarily co-expressed with other genes in the genetic circuits they are involved in, and so whether correlated expression domains indicate a related function or merely co-regulation by other factors. Genomic proximity to other genes of interest is another approach to deducing functional connections (Gil and Ulitsky 2020). However, lncRNAs produced from loci near other genes often do not appear to regulate their expression (Hezroni et al. 2020; Ramos et al. 2015). Here, we set out to explore the function of *Silc1* in the central nervous system and its relationship with *Sox11*.

## Results

### *Silc1* is broadly expressed in the central nervous system

In our initial description of *Silc1* we noted that in contrast to *Sox11*, which is expressed at higher levels at embryonic stages, *Silc1* was detected almost exclusively in postnatal samples of nervous systems and that in those postnatal samples *Silc1* expression was substantial and largely comparable to that of *Sox11* (Perry et al. 2018). We also noted that while *Silc1* was induced by ~10 fold following sciatic nerve injury, the RNA-seq–derived expression levels in the injured DRG were on par with its steady-state levels in the central nervous system, where it was not induced in various injury models (Perry et al. 2018). We, therefore first wanted to examine where in the postnatal brain *Silc1* and *Sox11* are expressed. We re-analyzed the RNA-seq data from the NeuroSeq atlas – a set of genetically-defined cellular populations isolated by a combination of fluorescent proteins and microdissection from the mature brain (Sugino et al. 2019) (**Fig. 1A**). Across the 517 samples in this dataset, *Sox11* was detected in 467 (90%) and *Silc1* in 504 (97%) with at least 5 reads per million in 40% of the samples for *Sox11* and 80% of the samples for *Silc1*, suggesting that both genes are rather broadly present in the mature brain. Across the samples, *Sox11* and *Silc1* exhibited a significant but overall modest correlation (Spearman R=0.25, P=2.46×10^-8^). *Sox11* expression was an order of magnitude higher in a single population in the hippocampus - the POMC-positive neuronal progenitors, where *Silc1* was barely detectable (RPKM<0.3, **Fig. 1A**), but both genes were expressed at similar and consistent levels of 3–5 RPKM across most other populations, with a notable lack of *Silc1* expression in the olfactory epithelium (**Fig. 1A**). Across all the mouse genes in the NeuroSeq dataset, *Silc1* was most closely correlated (R=0.57, P<10^-16^) with *Thy1*, a marker for neuronal maturation and cessation of neurite outgrowth (Bradley, Ramirez, and Hagood 2009), suggesting *Silc1* transcription switches on when these processes take place. Consistently with the RNA-seq data, fluorescence in situ hybridization (FISH) analysis using RNAscope as well traditional FISH showed broad expression of *Silc1* throughout the adult brain (**Fig. 1B**).

**Figure 1.**
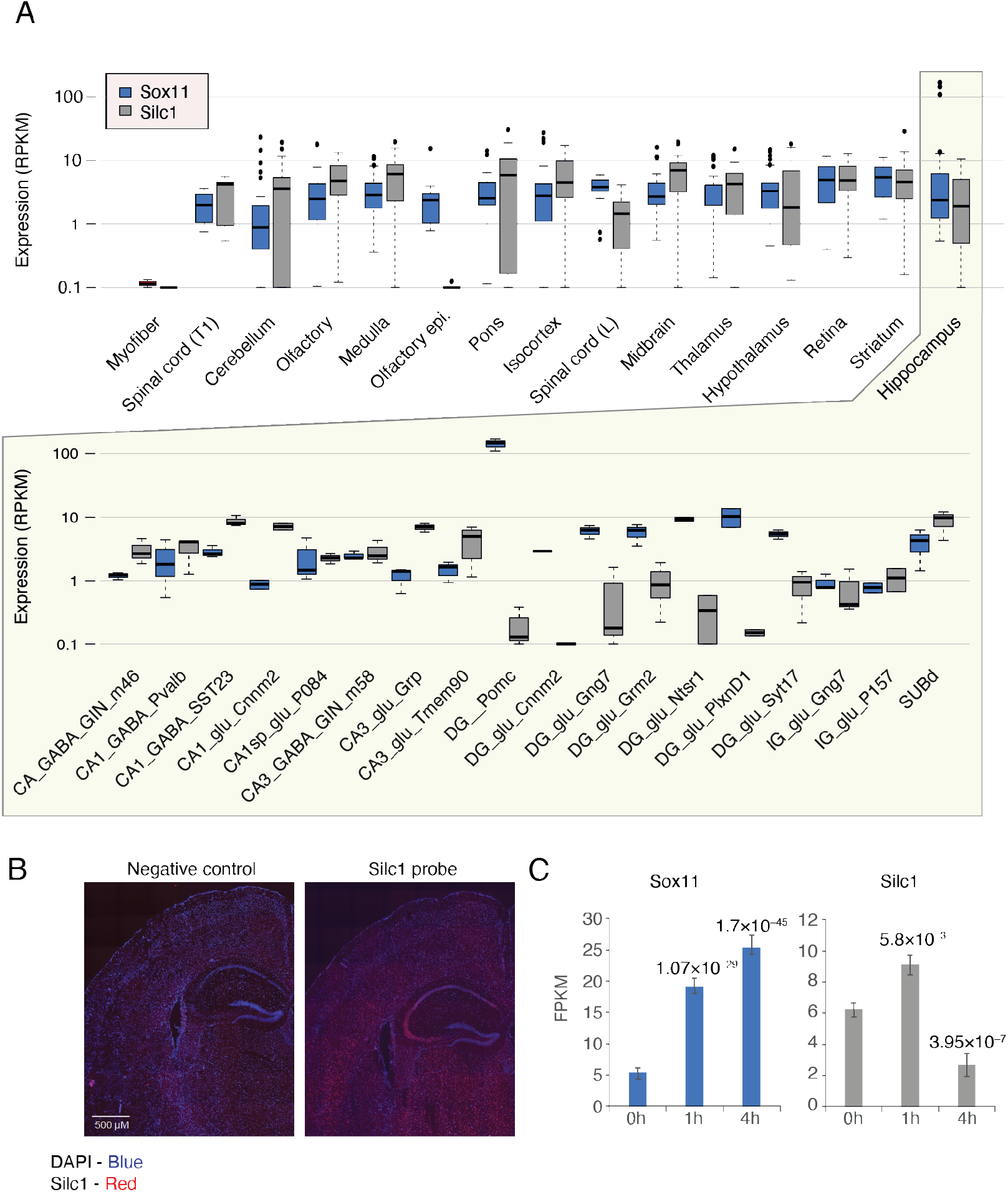
**(A)** Expression levels of the Sox11 (blue) and Silc1 (gray) in sorted populations from different regions of the CNS. Data from the Allen brain atlas. The bottom panel shows the different populations of cells within the hippocampus. **(B)** Fluorescent *in situ* hybridization (FISH) assay using RNAscope for brain sections from WT mouse. Negative control was performed in parallel as an indicator of background staining. Tissues were hybridized with *Silc1* probes (red) and counterstained with DAPI (blue), and imaged using 100X oil-immersion objective. Scale bar: 500 μm. **(C)** Quantification of the indicated genes in RNA-seq data from the DG at the indicated time points following electroconvulsive stimulation using RNA-seq data from (Su et al. 2017). P-values were computed for comparison with the ‘0h’ time point, using DeSeq2 (Love, Anders, and Huber 2014) and adjusted for multiple testing.

We conclude that a small population of neuronal progenitors in the hippocampus expresses exceptionally high levels of *Sox11* and no *Silc1*, closely resembling the embryonic progenitors. In other parts of the mature brain, both *Silc1* and *Sox11* are expressed, but with modestly correlated patterns that can be even anti-correlated when considering individual regions (see below).

**Figure S1.**
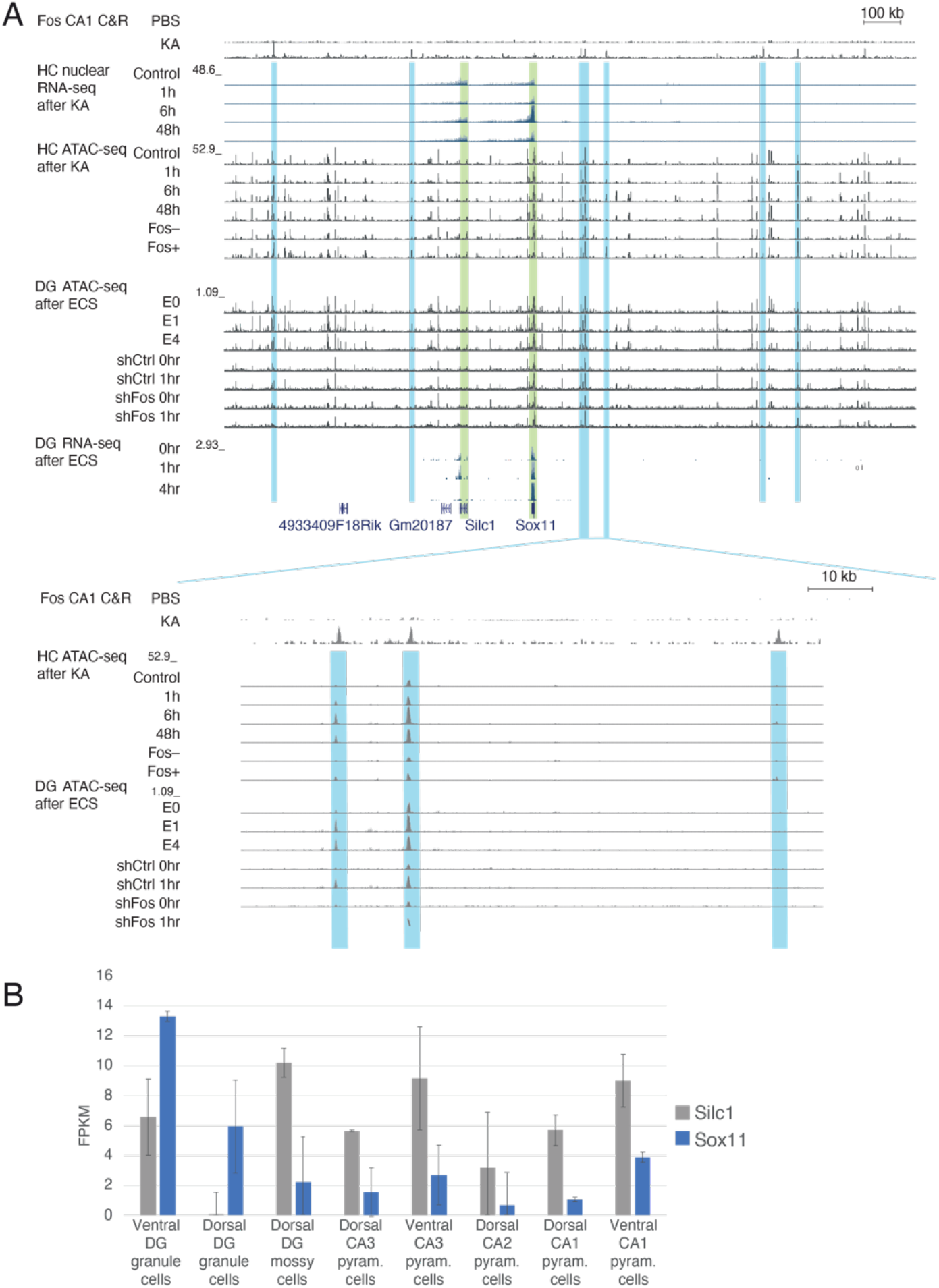
Transcriptomic and epigenomic read coverage in the broad Sox11 domain. **(A)** Top: broad ~2Mb region flanking Sox11; Bottom: zoom-in on the indicated region. Shown are (top to bottom): Fos Cut&Run data in the hippocampus (HC) CA1 region (Fernandez-Albert et al. 2019); HC nuclear RNA-seq data at the indicated time after KA treatment (Fernandez-Albert et al. 2019); ATAC-seq data from the same study; including from sorted Fos-positive and Fos-negative cells; ATAC-seq data from the DG after ECS (Su et al. 2017), including a time course and the Fos knockdown experiment; RNA-seq data from the same study. Fos-bound and apparently Fos-regulated regions are shaded. **(B)** Expression of *Sox11* and *Silc1* in data of sorted populations from the HippoSeq dataset.

### *Silc1* and *Sox11* are differentially induced by neuronal activity in the hippocampus

To focus on a particular region in the adult brain where *Silc1* may play a relevant role, we examined public RNA-seq data and literature for evidence of changes in the expression of *Silc1* and/or *Sox11* in different physiological settings. We noted that upon electroconvulsive stimulation, *Sox11* mRNA was previously shown to be specifically induced in the DG (von Wittgenstein et al. 2020; Sun et al. 2005), and examined RNA-seq–based expression in the DG upon stimulation, which showed that both *Silc1* and *Sox11* are induced at early time points, followed by a decline of *Silc1* to significantly below-basal levels (**Fig. 1C**). We further examined ATAC-seq data from the same study, and from the hippocampus of mice stimulated with kainic-acid (Fernandez-Albert et al. 2019) and found several specific activity-induced enhancers in the gene deserts flanking *Sox11*, which also overlapped binding sites for the AP-1 complex (**Fig. S1A**), suggesting a plausible regulatory route for neural activity-driven induction of the two genes.

Interestingly, when zooming into the hippocampus cell populations of the NeuroSeq dataset, *Silc1* and *Sox11* expression patterns were strongly anticorrelated (Spearman R=–0.45, P=8.2×10^-4^, **Fig. 1A**), with *Sox11* mRNA predominantly expressed in the DG and *Silc1* in the Amonion Horn (AH) sub-regions. Similar results were observed in a HippoSeq dataset (**Fig. S1B**) (Cembrowski et al. 2016). Therefore, while on the tissue level, both *Sox11* and *Silc1* appear to be regulated by neuronal activation, similar to their co-induction in the DRG (Perry et al. 2018), the baseline expression patterns of the two genes in the hippocampus appear to be strikingly different.

### *Silc1* is required for efficient memory formation

Since the hippocampus is known to be associated with spatial memory acquisition, we then turned to examine the consequences of lack of *Silc1* on memory formation, using two commonly used paradigms for spatial learning, the Morris water maze (MWM) and the Barnes maze (Matthew W. Pitts, 2018). MWM training consisted of 8 daily sessions, each comprised of four trials (from different starting points). In each training trial, mice were allowed to swim either until they located the platform or until 90 seconds elapsed. The latency (sec) to reach the platform was recorded. Whereas among WT mice the latency to find the platform was significantly reduced after 4 daily sessions, *Silc1^-/-^* mice exhibited impaired learning, as their latency to reach the platform was significantly reduced only after 6 days **(Fig. 2A).** In the probe trial (memory recall test) that was performed 24 hours following the last training session, no significant differences were observed between the genotypes in either index (quadrants total distance and cumulative duration) **(Fig. 2B).** A similar pattern of results was also observed in the Barnes maze – WT and *Silc1^-/-^* mice were trained for a total of four days, receiving four trials every day; latency (sec) to enter the escape tunnel was recorded. Significant differences were observed between the genotypes only during learning but not during the probe trial (**Fig. 2C-D).** Finally, mice underwent Fear Conditioning, and were tested for both hippocampal-dependent, contextual memory as well as amygdala-dependent, cue memory (**Fig. S2**). No differences were noted between the genotypes during the conditioning phase or either of the memory tests.

**Figure 2.**
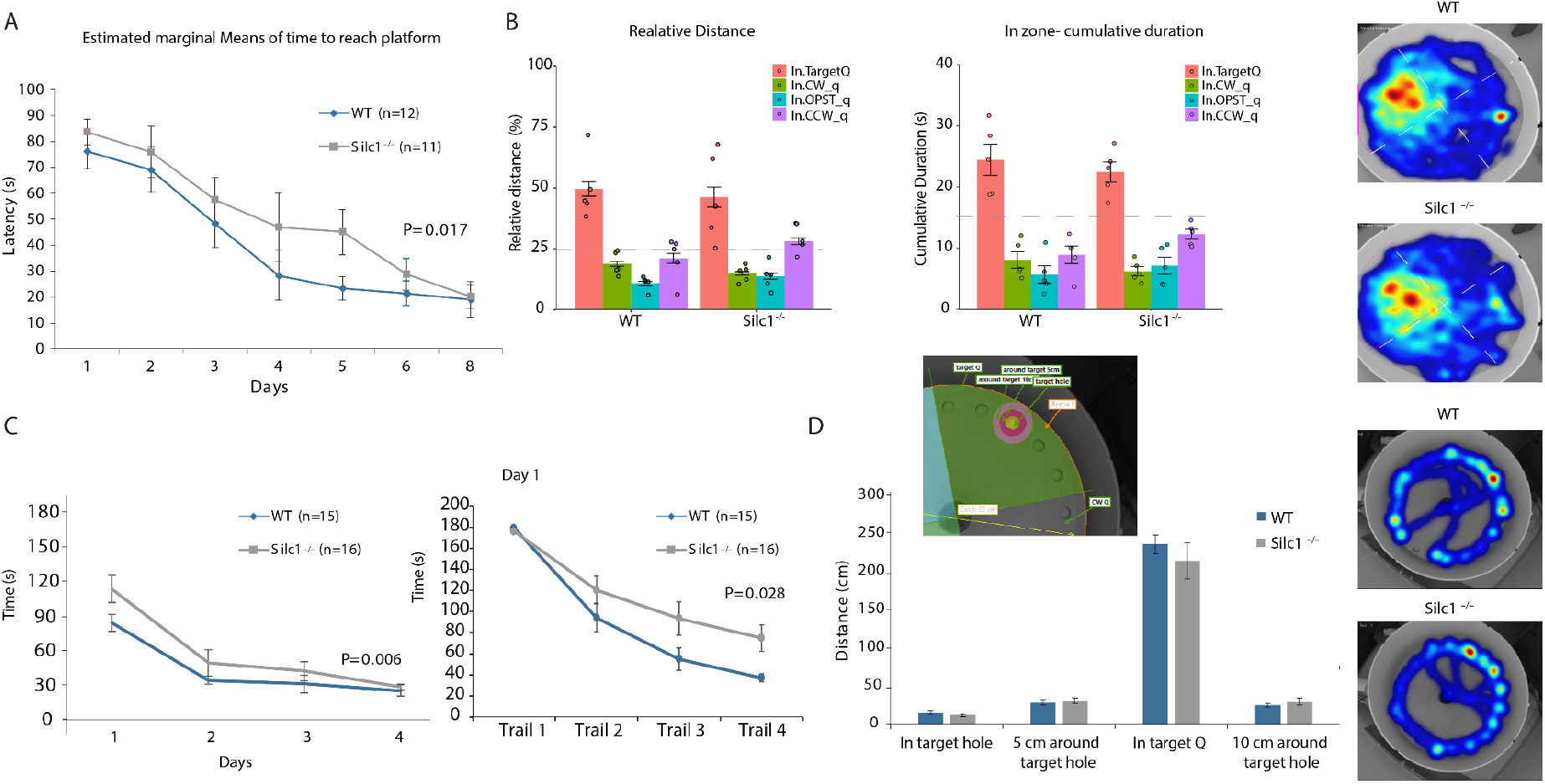
*Silc1^-/-^* mice showed slower spatial learning in the Morris water maze and Barnes maze. **(A)** Escape latencies (sec.) over MWM training sessions; *Silc1^-/-^* mice exhibited significantly slower learning. **(B)** Probe test session (24 hr after acquisition); total distance swam (cm) and cumulative duration (sec) in each of the quadrants. Grouped heatmaps of cumulative duration during the probe test indicate that *Silc1^-/-^* and WT mice explored the target quadrant similarly. **(C)** Escape latencies (sec) in the Barnes maze over daily training sessions (left) and with-in trials of Day 1 (right); *Silc1^-/-^* mice exhibited significantly slower learning. **(D)** Probe test session (24 hr after last training session); total distance walked (cm) in each of the quadrants during the probe test. Grouped heatmaps of cumulative duration during the probe test indicate that *Silc1^-/-^* and WT mice explored the escape hole target quadrant similarly. Data represent mean +/- SEM (error bars). Two-way ANOVA, for Gene (Between-Subjects), Training Days (Within-Subjects with repeated measures), and their interaction (Gene × Training Days); followed by planned contrasts analyses (repeated measures) used to measure the significance of the effect on training.

**Figure S2.**
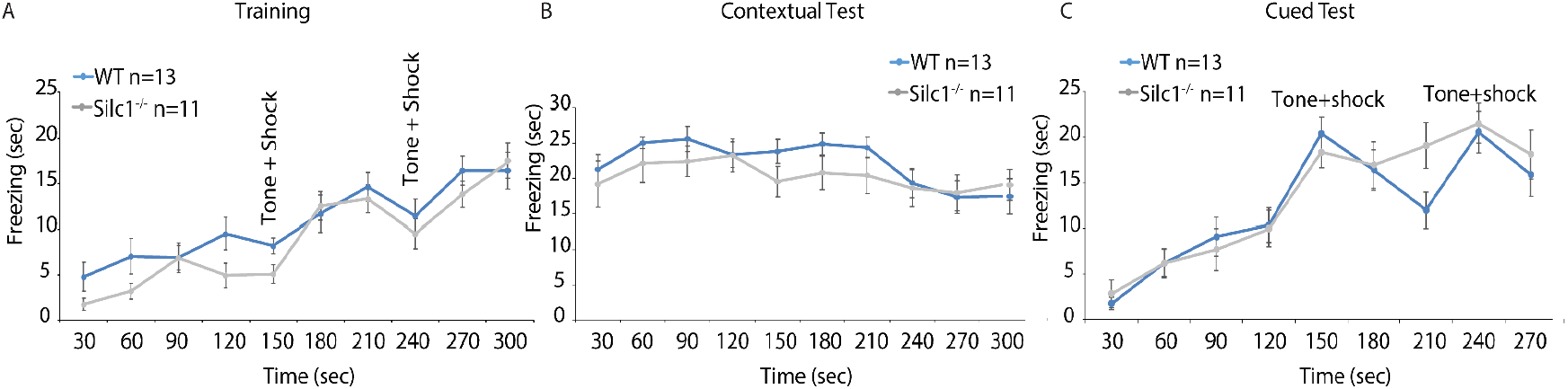
*Silc1^-/-^* mice showed no alteration in freezing responses during fear conditioning (A) and subsequent recall tests. Panels A, B and C show the absence of differences in the fear conditioning freezing behavior of *Silc1^-/-^* mice compared to WT littermates. Data represent mean +/- SEM (error bars). Two-way ANOVA, for Gene (Between-Subjects), Time (30 sec intervals; Within-Subjects with repeated measures), and their interaction (Gene × Time).

### *Silc1* and *Sox11* are immediate early genes up-regulated in mice placed in a novel environment

Since an intact *Silc1* locus was associated with a response of mice to a novel stimulus, we then used the exposure to the Barnes maze setting at several time points (0.5, 1, 2, and 6 hours) as a ‘novel environment’ paradigm (NE). We first used RNAscope to map the expression of *Silc1* and *Sox11* in the home cage (HC) and the NE. We observed the highest induction of *Silc1* at 1 hr post-NE **(Fig. S3A)**, similar to the response of other immediate early genes like *Fos* and *Arc*, and continued to use this setting in subsequent experiments. As it has been reported that the *Sox11* coding sequence (CDS) and 3’ UTR are not always observed in strictly the same cells or regions (Struebing et al. 2017), we used separate sets of probes targeting the CDS and the 3’ UTR. Consistent with the data from the FACS-sorted populations described above, in control conditions *Silc1* was higher in the AH than in the DG, whereas *Sox11* mRNA was more abundant in the DG. In NE, the expression of both RNAs increased in both regions **(Fig. 3 and S3A)**. When examining SOX11 protein, surprisingly, the most notable increase was in the AH region **(Fig. 4A)**. We validated the specificity of this signal by Cre injection into the hippocampus of *Sox11*^fl/fl^ mice from the Lefebvre lab (Bhattaram et al. 2010) (**Fig. 4B and 4D**). Strikingly, in *Silc1^-/-^* mice, there was a significant reduction in *Sox11* mRNA and protein levels in both the HC and NE conditions that was noted in both the DG and AH regions **(Fig 3B and 4A)**. Consistently with previous reports, *Sox11* CDS and 3’UTR expression patterns did not always strictly overlap, and loss of *Silc1* resulted in a concordant decrease in both signals in the DG and CA3 regions (**Fig. 3B**). Furthermore, we observed a substantial reduction in the protein expression of SOX11 in the AH. These results were verified by a Western blot analysis with SOX11 antibody using proteins extracted from the hippocampus of WT and *Silc1^-/-^* mice under HC and NE conditions. SOX11 protein levels were significantly reduced in *Silc1^-/-^* mice upon stimulus in the NE setting (**Fig. 4D-E**). Assessing the colocalization of Fos-positive cells and Sox11 in the DG and CA3 regions indicated low rates of cells with prominent colocalization (**Fig. S4**), suggesting that the *Sox11* expression is not limited to the Fos-positive cells that presumably experienced the strongest activation.

**Figure 3.**
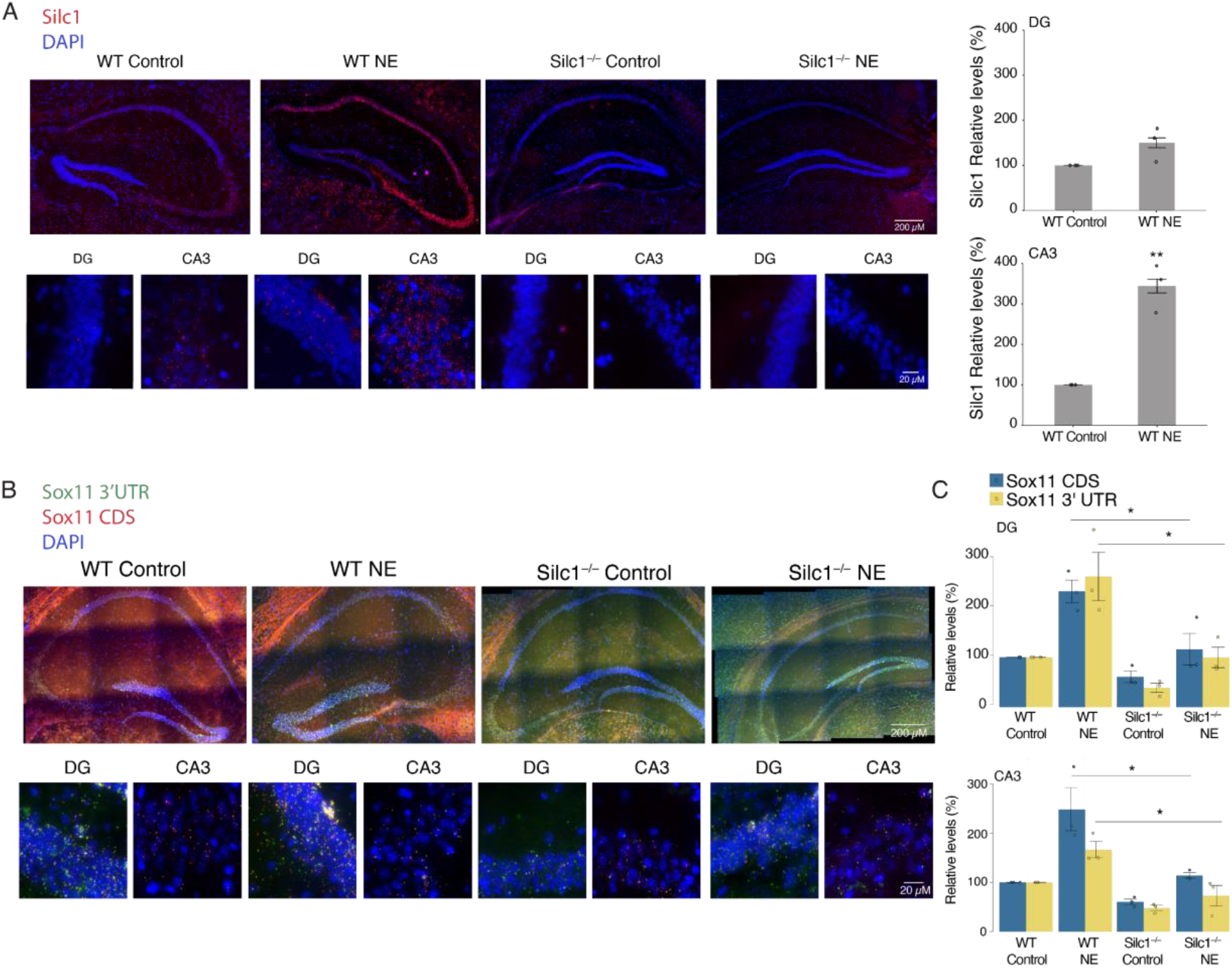
Expression of *Silc1* and *Sox11* in control and novel environment (NE) conditions in WT and *Silc1^-/-^* mice. **(A)** RNAscope fluorescent *in situ* hybridization (FISH) assay on hippocampal sections from mice with the indicated genotype. Tissues were counterstained with *Silc1* probe (red) and DAPI (blue) and imaged using 20X (Scale bar 200 μm) and 100X oil-immersion objectives (Scale bar 20 μm). Novel environment (NE) was performed using the Barnes maze setting, and the hippocampus was extracted for coronal sections after 1 hr of stimulus. 12 images of non-overlapping fields per biological repeat were quantified, 3 biological repeats. Mean ± SEM is shown. P value calculated using unpaired two sample t-test, ** P < 0.005. **(B)** As in A with tissues hybridized with *Sox11* CDS (red), 3’UTR (green) probes and counterstained with DAPI (blue). **(C)** As in A for the number of green and red dots. P value calculated using unpaired two sample t-test, * P < 0.05.

**Figure 4.**
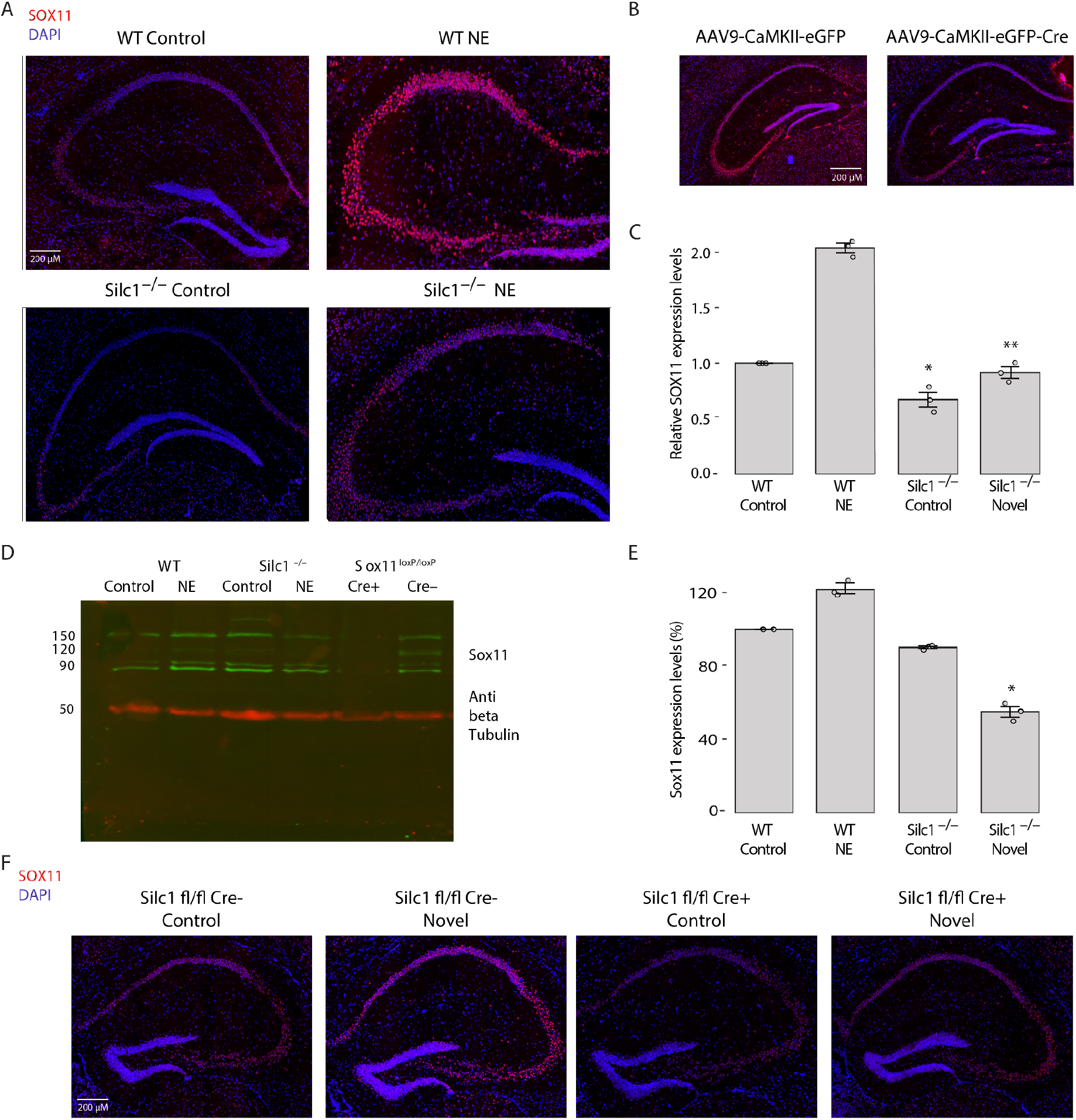
Levels of SOX11 protein following novel environment in WT and Silc1^-/-^ hippocampus. **(A)** Immunostaining with anti-SOX11 (red) and DAPI (blue) in hippocampi of WT and *Silc1^-/-^* mice in HC and NE conditions. Imaging using 20X objective (Scale bar 200 μm). **(B)** As in A for Sox11^fl/fl^ mice injected stereotaxically in the CA3 region with Cre-GFP or GFP-expressing AAV9 viruses. Imaging was done using 20X objective (Scale bar 200 μm). SOX11 levels were significantly reduced after Cre injection, which indicates the specificity of the SOX11 antibody used for staining. **(C)** Quantification of 3 biological repeats of hippocampal staining. Mean ± SEM, * P < 0.05, ** P < 0.005, unpaired two-sample t-test. **(D)** Western blot with SOX11 and beta-tubulin antibodies from mice with the indicated genotype, using whole-hippocampal protein extract. Hippocampus protein extract from Sox11^fl/fl^ mice that were stereotaxically injected in the CA3 region using AAV9 Cre-GFP or AAV9 GFP were used as a specificity control for the antibodies. **(E)** Western blot quantification, SOX11 expression levels were normalized to beta-tubulin levels. n=3, Mean ± SEM is shown, * p < 0.05, unpaired two-sample t-test. **(F)** As in A for hippocampus sections from *Silc1^-/-^* mice that were stereotaxically injected in the CA3 region using AAV9 Cre-GFP or AAV9 GFP. Two weeks after injections, the mice were exposed to a novel environment (NE) and the hippocampus was extracted for coronal sections after 1 hr of NE. Imaging using 20X objective (Scale bar 200 μm).

### Conditional depletion of *Silc1* in the adult brain leads to reduced SOX11 expression

*Silc1^-/-^* mice lack the *Silc1* promoter and never express *Silc1*, so the observations of changes in SOX11 in the *Silc1^-/-^* mice may reflect changes in embryonic brain development or postnatal brain maturation. To address this, we generated Silc1 conditional mice by inserting loxP sites at regions flanking the *Silc1* promoter (using the same CRISPR gRNAs used for generation of *Silc1^-/-^* mice).

For specific *Silc1* reduction in the adult hippocampus, AAV9 Cre-GFP or AAV9 GFP were injected into the CA3 region of the hippocampus of *Silc1^fl/fl^* mice. A local reduction in *Silc1* levels was detected only around the Cre injection site (**Fig. S3B**). In addition, SOX11 protein levels were assessed using immunostaining and indicated a specific reduction in SOX11, following Cre injection, only in cells that expressed GFP **(Fig. 4F).** Taken together, the findings demonstrated that *Silc1* lncRNA, *Sox11* mRNA, and SOX11 protein are all induced in the CA3 region when mice are exposed to NE, and *Silc1* depletion leads to a reduction of Sox11 mRNA and protein in the hippocampus.

**Figure S3.**
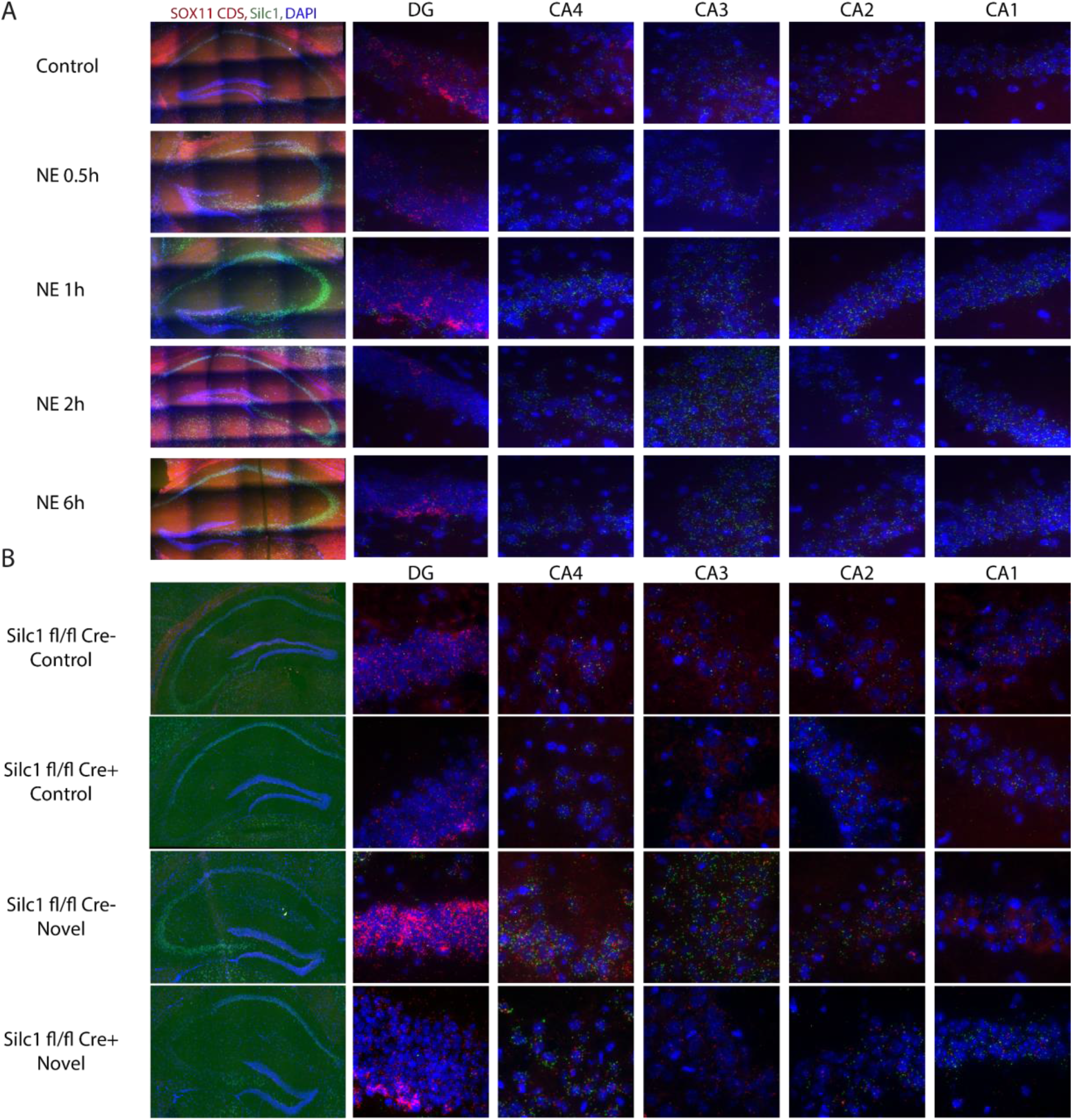
Expression of *Silc1* and *Sox11* at several time points following novel environment (NE) conditions in *Silc1* conditional knockout mice. **(A)** RNAscope FISH assay on hippocampal sections from WT mice. Tissues were hybridized with *Silc1* (green) and Sox11 CDS (red) probes and counterstained with DAPI (blue), and imaged using 20X (Scale bar 200 μm) and 100X oil-immersion objectives (Scale bar 20 μm). Novel environment (NE) performed using the Barnes maze setting and the hippocampus was extracted for coronal sections 0.5, 1, 2 and 6 hr upon stimulus. **(B)** As in A for hippocampus sections from *Silc1^fl/fl^* mice that were stereotaxic injected in the CA3 region using AAV9 Cre-GFP or AAV9 GFP. Two weeks after injections the mice were exposed to Novel environment (NE) and the hippocampus was extracted for coronal sections after 1 hr of NE.

**Figure S4.**
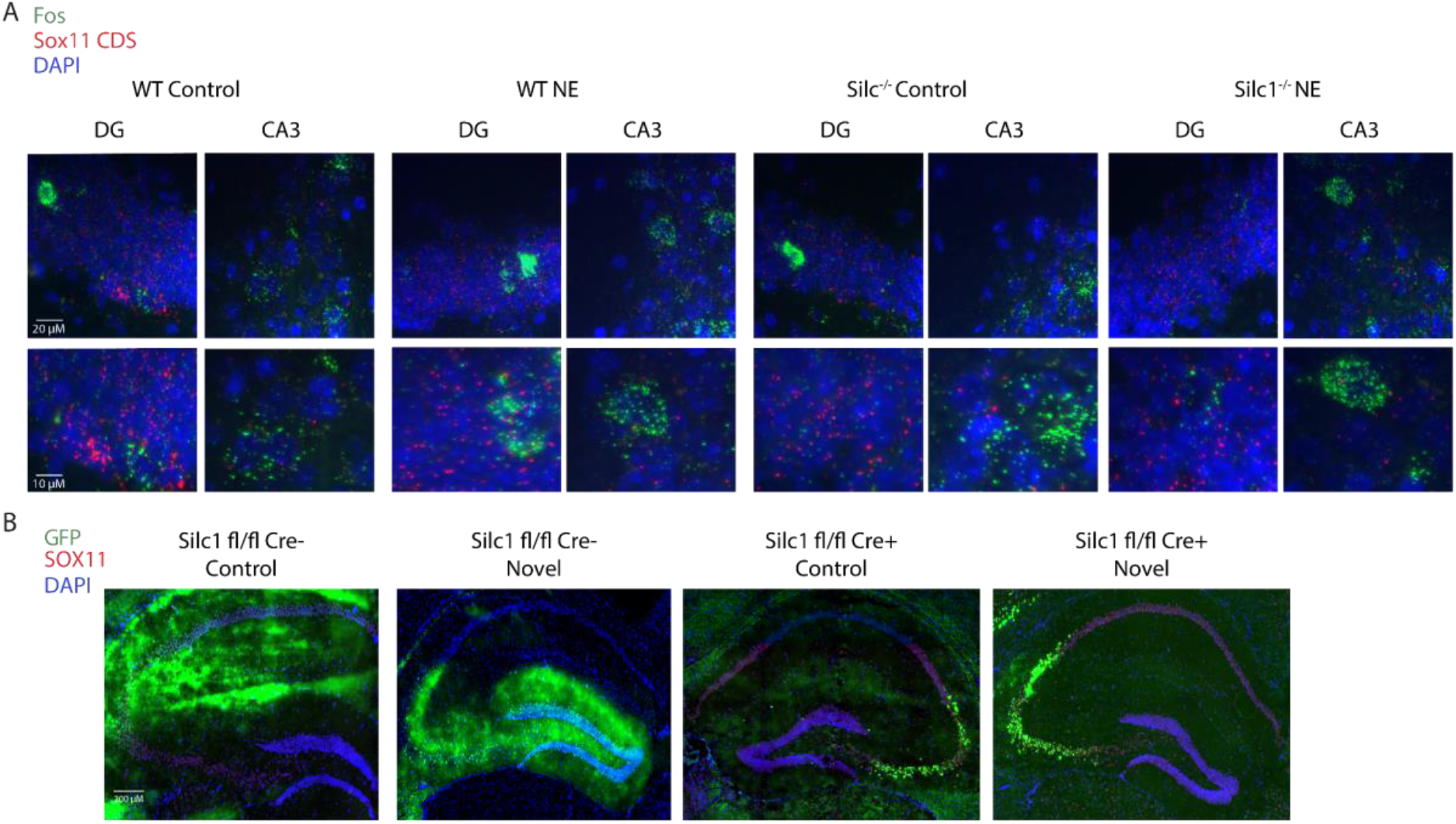
Relationship between cells with *Sox11* RNA expression and Fos-positive cells. **(A)** RNAscope Fluorescent *in situ* hybridization (FISH) assay on WT and *Silc1^-/-^* mice hippocampal sections from HC and NE conditions. Tissues were hybridized with Fos mRNA (green) and Sox11 CDS (red) probes and counterstained with DAPI (blue), and imaged using 100X oil-immersion objectives (Scale bar 20 μm). **(B)** Immunostaining with anti-SOX11 (red) and DAPI (blue) in hippocampi of *Silc1^fl/fl^* mice that were stereotaxically injected in the CA3 region using AAV9 Cre-GFP or AAV9 GFP. The GFP signal marks the site of injection. Imaging using 20X objective (Scale bar 200 μm).

### Knockdown of *Silc1* in the adult hippocampus

When using the *Silc1^-/-^* or *Silc1^fl/fl^* mice it is impossible to test separately the relative contribution of the *Silc1* promoter, which they are lacking, and the role of the transcription of *Silc1* and/or its RNA product. To tease apart these effects, we first attempted to use CRISPR/Cas9 to introduce a polyadenylation signal into the first exon of *Silc1*, but this only partially reduced *Silc1* expression in animals carrying homozygous insertions (**Fig. S5A**). Therefore, we opted for using GapmeRs – antisense nucleotides – to degrade the *Silc1* RNA product, potentially affecting also its transcription without altering the DNA of the locus (Lee and Mendell 2020; Lai et al. 2020). We first used a primary DRG culture to select GapmeRs that effectively reduce the expression levels of *Silc1* or *Sox11* (with separate GapmeRs targeting *Sox11* CDS and its 3’ UTR). Efficient GapmeRs were selected by transfection into cultured DRG neurons (**Fig. S5B**). We then introduced these GapmeRs specifically into the CA3 region of the hippocampus by stereotaxic injection and assessed the expression of *Silc1* and *Sox11* by RNAscope. Five days after injection of the *Silc1*-targeting GapmeR, we observed a local reduction in the expression of *Silc1*, which coincided with reduced levels of the CDS and 3’UTR of *Sox11*, as well as of SOX11 protein (**Fig. 5A-D**). GapmeRs targeting *Sox11*, on the other hand, reduced *Sox11* levels (further confirming the specificity of our detection reagents) but did not notably affect the expression of *Silc1* (**Fig. 5E-H and Fig. S6**). These results indicate that transcription through the *Silc1* locus or the *Silc1* RNA product in the mature adult neurons are essential for the regulation of *Sox11* levels.

**Figure 5.**
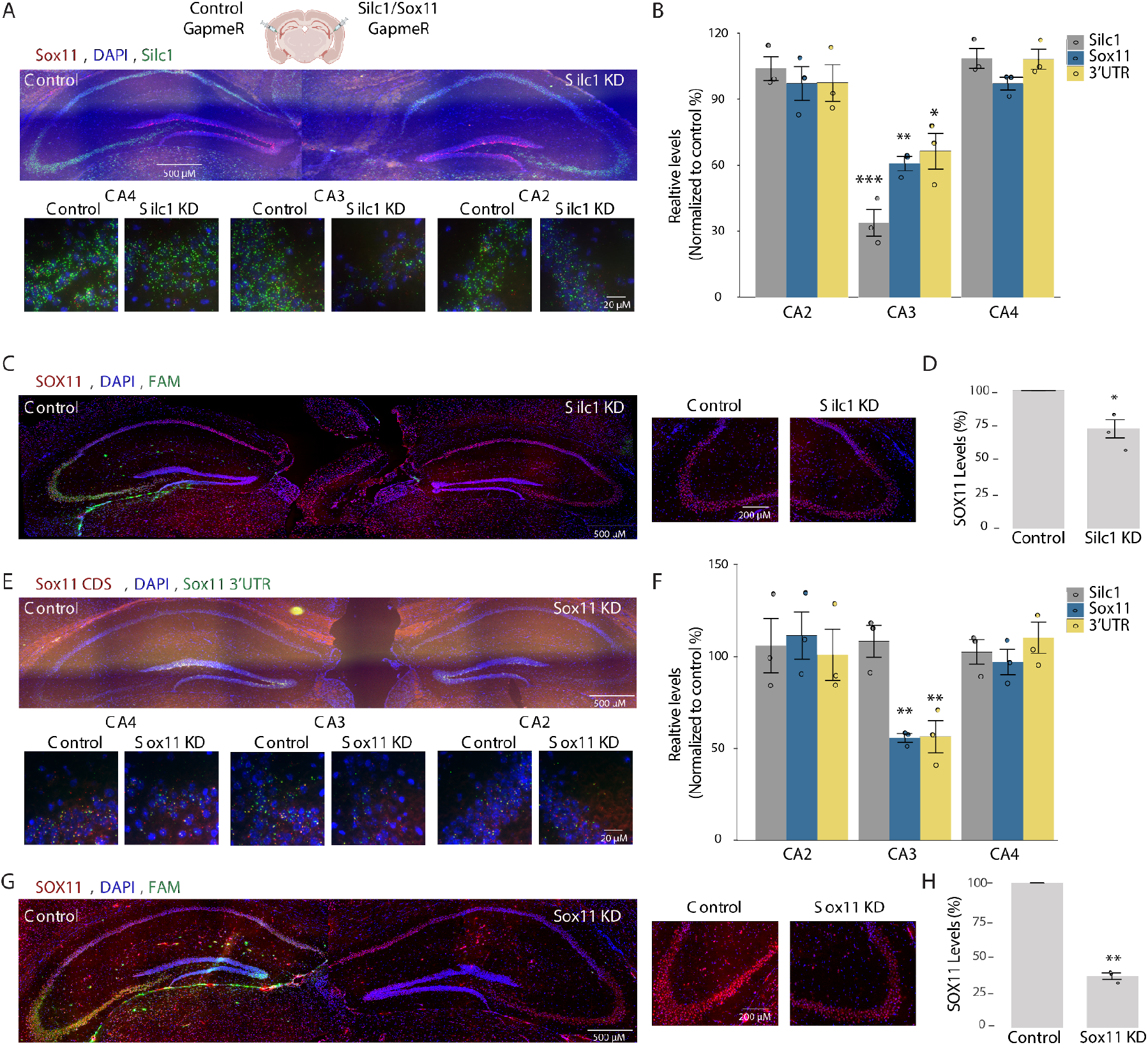
*Silc1* and *Sox11* KD by injections of GapmeRs to the CA3 region. **(A)** Control FAM-labelled (left) or *Silc1*-targeting (right) GapmeRs were injected into the CA3 region. 5 days later, the hippocampus was extracted for coronal sections. RNAscope analysis of *Silc1* and *Sox11* expression using Silc1 (green), Sox11 CDS (red) probes, and DAPI. Imaging was done using 20X (Scale bar 200 μm) and 100X oil-immersion objectives (Scale bar 20 μm). **(B)** RNAscope quantification of the number of green and red dots, normalized to control GapmeR, performed using IMARIS software. 12 images of non-overlapping fields were quantified per biological repeat; 3 biological repeats. Mean ± SEM is shown. P value calculated using an unpaired two-sample t-test, * P < 0.05, ** P<0.005, *** P<0.001. **(C)** Immunostaining with anti-SOX11 (red) and DAPI (blue) in hippocampi of *Silc1* KD mice. FAM signal marks the injection site of the control GapmeR. Imaging using 20X objective (Scale bar 200 μm). **(D)** Quantification of 3 biological repeats of hippocampal staining, normalized to injection of control GapmeR. Mean ± SEM, * P < 0.05, unpaired two-sample t-test. **(E)** As in A using Control (left) or Sox11 CDS (right) GapmeRs. **(F)** As in B using Control or Sox11 CDS GapmeRs. **(G)** As in C for *Sox11* KD mice. **(H)** As in D for *Sox11* KD mice.

**Figure S5.**
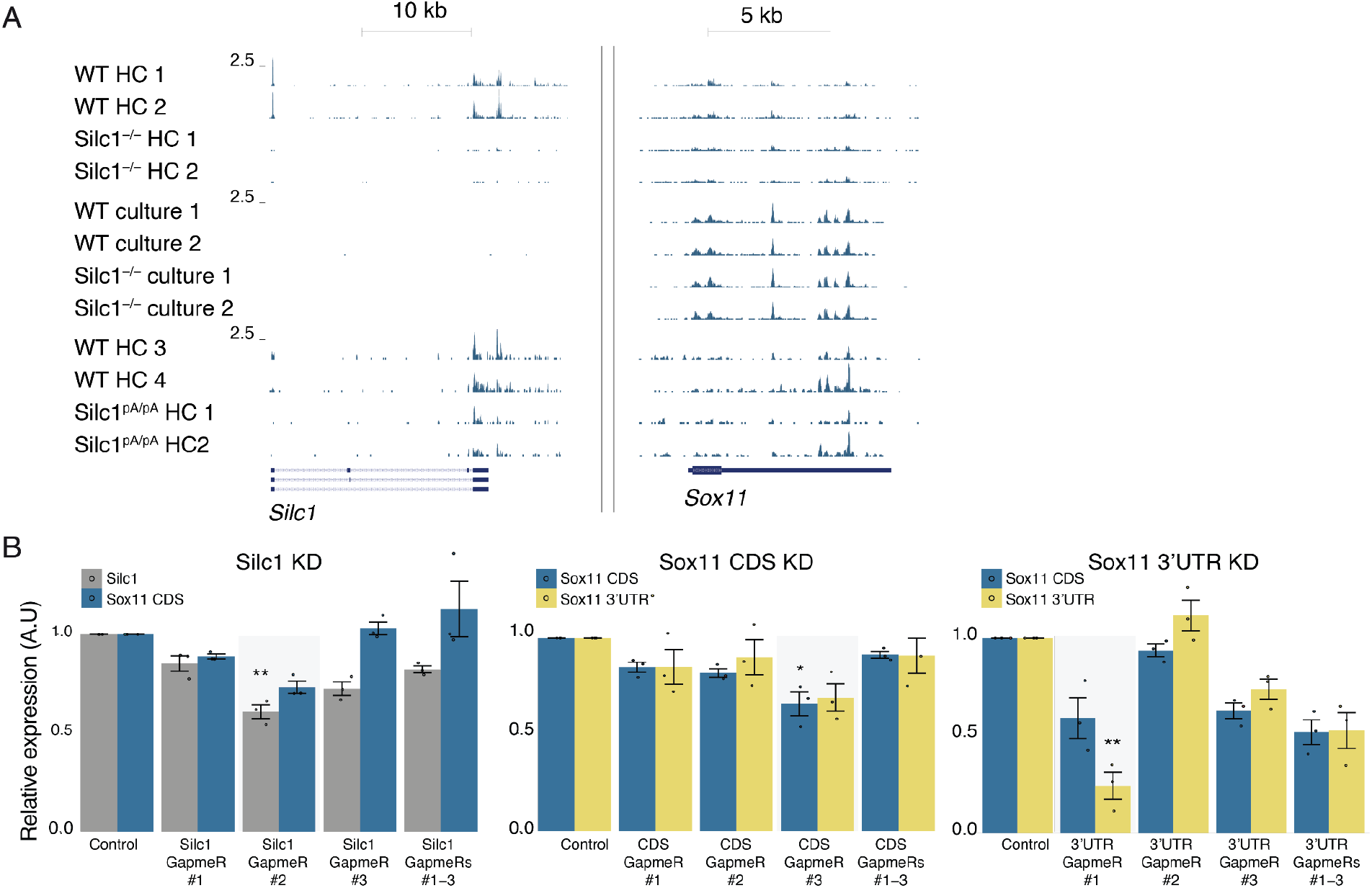
Methods to perturb and study *Silc1* transcription. (A) RNA-seq read coverage at the *Silc1* (left) and *Sox11* (right) genomic regions, in the adult hippocampus (HC) or in cultured hippocampal neurons (“culture”) from the indicated genetic background. (B) Changes in expression of the indicated genes and regions upon the use of the indicated GapmeRs targeting *Silc1* (left) *Sox11* CDS (middle) and *Sox11* 3’UTR (right), as evaluated by qRT-PCR. Levels were normalized to control GapmeR and β-actin for internal control. The GapmeRs selected for further experiments are shaded in gray.

**Figure S6.**
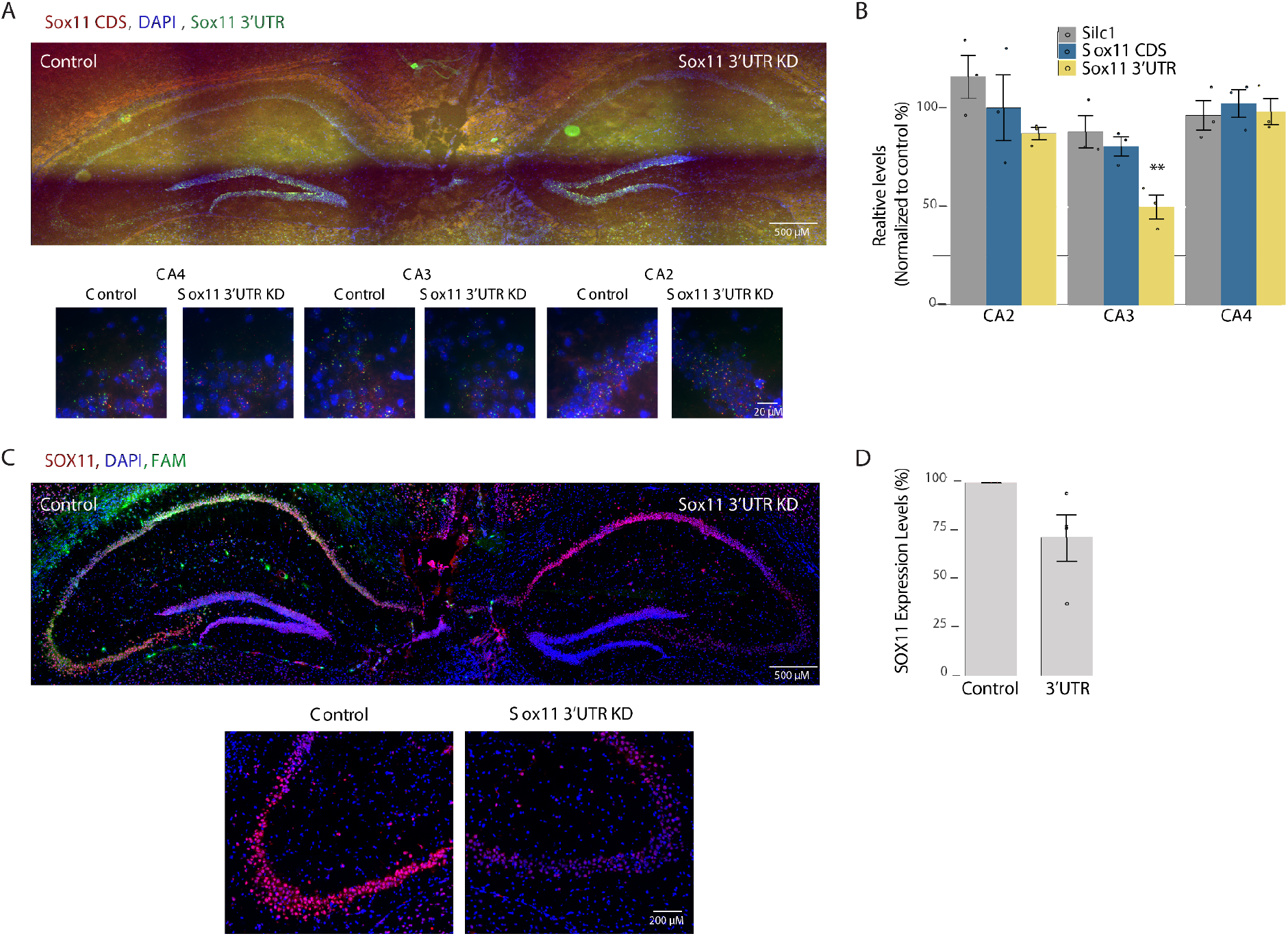
Expression of *Sox11* after *Sox11* 3’ UTR KD by injection of GapmeRs into the CA3 region. **(A)** Control-FAM (left) or Sox11 3’ UTR (right) GapmeRs were injected into the CA3 region. 5 days later, the hippocampus was extracted for coronal sections. RNAscope analysis of *Sox11* expression using Sox11 3’ UTR(green), Sox11 CDS (red) probes, and DAPI. Imaging was done using 20X (Scale bar 200 μm) and 100X oil-immersion objectives (Scale bar 20 μm). **(B)** RNAscope quantification of the number of green and red dots, normalized to control GapmeR, performed using IMARIS software. 12 images of nonoverlapping fields per biological repeat were quantified; 3 biological repeats. Mean ± SEM is shown. P value calculated using an unpaired two-sample t-test, * P < 0.05. **(C)** Immunostaining with anti-SOX11 (red) and DAPI (blue) in hippocampi of *Sox11* 3’ UTR KD mice. FAM signal marks the injection site of the control GapmeR. Imaging using 20X objective (Scale bar 200 μm). **(D)** Quantification of 3 biological repeats of hippocampal staining, normalized to the injection of control GapmeR. Mean ± SEM.

### Overexpression of the *Silc1* RNA sequence does not affect *Sox11* or downstream genes

In the DRG, *Silc1* regulates *Sox11* expression strictly in *cis*, and over-expression of Silc1 cDNA did not alter *Sox11* levels (Perry et al., 2018). Therefore, we assessed whether that is the case also in the hippocampus. In addition, we sought a signature for changes in gene expression following the up-regulation of *Sox11* in the AH, where it has not been studied to date. To that end, we used AAV9 injected into the CA3 region to over-express SOX11. The AAV vector expresses a GFP mRNA either as a control or fused to the Sox11 coding sequence, or the Silc1 sequence, under the CaMKIIα promoter specific to neurons (X. Wang et al. 2013). As depicted in **Fig. 6A-B**, two or three weeks after introduction, the GFP signal diffused to other regions of the hippocampus, and we observed a significant increase in the expression of *Silc1* or *Sox11* but no substantial or significant cross-regulation between the two genes (**Fig. 6A-B**). *Silc1* overexpression did not affect SOX11 protein levels **(Fig. 6C)**. We then injected AAV9 viruses overexpressing Silc1 into the CA3 region of *Silc1^-/-^* mice. *Silc1* levels were rescued but without any notable effect on *Sox11* transcript or protein levels (**Fig. S7**). These data suggest that in as in the DRG, the overall presence of the *Silc1* RNA sequence in the cell does not affect *Sox11* levels, moreover, the function of the *Silc1* RNA product, if any, is only relevant when it is produced from its endogenous locus.

**Figure 6.**
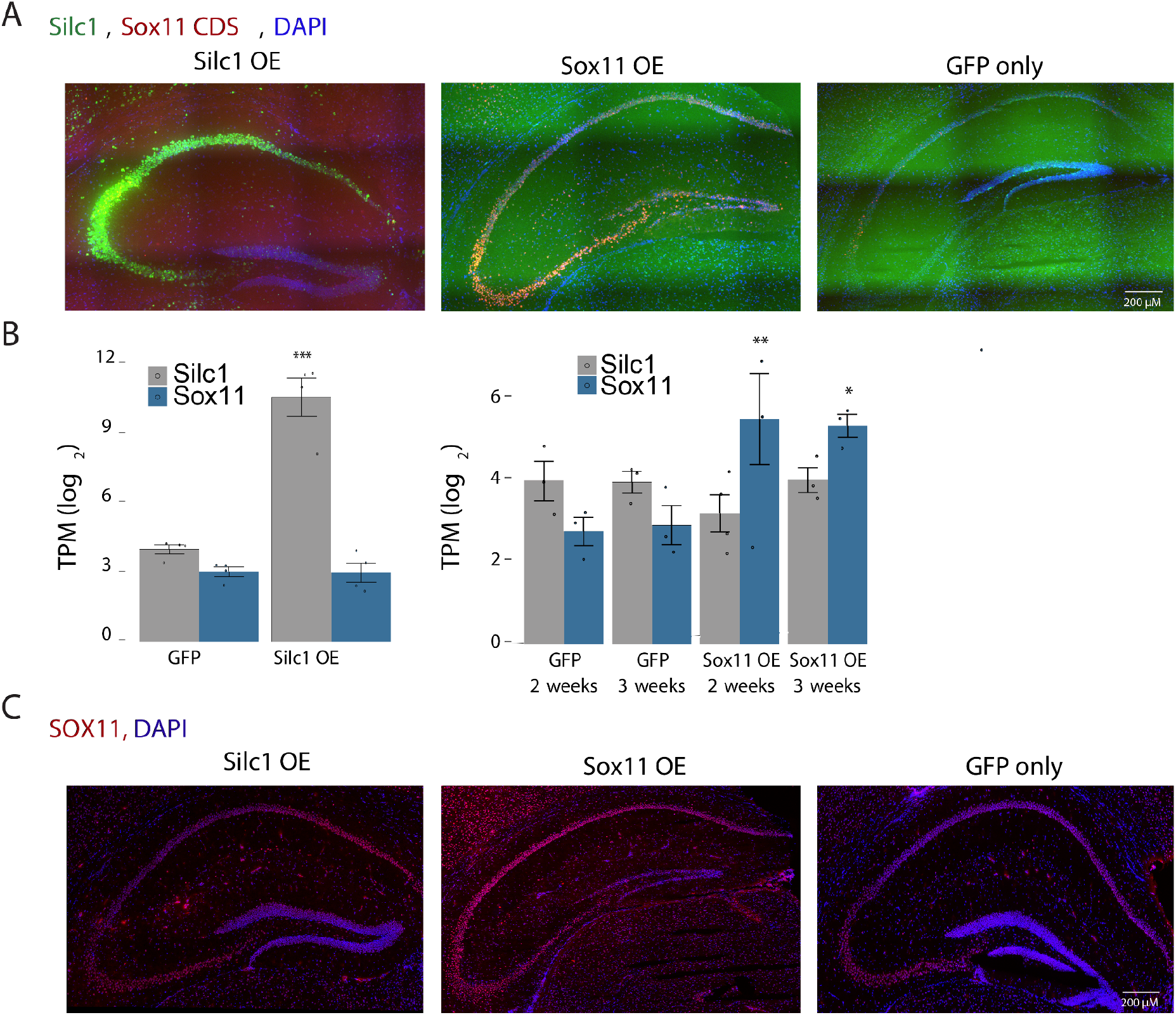
Overexpression of *Sox11* and *Silc1* in the hippocampus. **(A)** AAV9-Silc1, AAV9-Sox11 or AAV9-GFP were injected to the CA3 region for *Silc1* or *Sox11* overexpression. After 2-3 weeks the hippocampus was extracted for coronal sections. RNAscope analysis of *Silc1* and *Sox11* expression using Silc1 (green), Sox11 CDS (red) probes, and DAPI. Imaging was done using 20X (Scale bar 200 μm) and 100X oil-immersion objectives (Scale bar 20 μm). **(B)** RNA-seq quantification of *Silc1* and *Sox11;* 3 biological repeats. Mean ± SEM is shown. P value calculated using unpaired two-sample t-test, * P < 0.05,** P < 0.005, *** P < 0.001. **(C)** Immunostaining with anti-SOX11 (red) and DAPI (blue) in hippocampi of *Silc1* and *Sox11* OE mice. Imaging using 20X objective (Scale bar 200 μm).

### SOX11 drives a specific gene expression program in the adult hippocampus, which is affected by the loss of *Silc1*

While the Sox11-driven transcriptional program has been studied extensively in other settings, it is not known which transcription is sensitive to Sox11 levels in the mature adult neurons. To characterize these changes, we used RNA-seq to measure gene expression in the hippocampus of *Sox11*^flox/flox^ mice injected with an AAV vector driving expression of Cre (or GFP control) as compared with WT mice hippocampus over-expressing (OE) *Sox11* fused with GFP (or GFP control). Two or three weeks post-injection, the hippocampus was extracted and subjected to RNA-seq. Three weeks after the injection, we observed a reduction in both GFP and *Sox11* transcript levels, possibly due to cell toxicity caused by SOX11 overexpression (as observed in other systems (Norsworthy et al. 2017)) or silencing of the transgene (**Fig 6B**). In the RNA-seq data after 3 weeks of SOX11 OE, the 144 significantly reduced genes (fold-change <0.5 and adjusted P<0.05) were significantly enriched with genes down-regulated in neurodegenerative diseases: Huntington’s disease (Enrichr analysis, adjusted P=1×10^-13^) and Parkinson’s disease (P=2.85×10^-8^).

Therefore, focused on the RNA-seq data from the hippocampus extracts that were collected 2 weeks after AAV introduction for OE or one week after Cre injections. Among the 14,181 expressed genes (average FPKM>=.5, **Table S1**), we further focused on genes that changed significantly (P<0.05) in opposite directions by at least 50%. Under these definitions, 33 genes were positively regulated by Sox11 in the adult hippocampus (**Fig. 7A**), including the well-characterized targets *Dcx* (Haslinger et al. 2009) and *Mex3a* (Oliemuller et al. 2020), and only 7 were negatively regulated. These data are consistent with SOX11 acting predominantly as an activator, and therefore, we focused on the positively-regulated genes in further analyses. These genes were also significantly up-regulated in published datasets of AAV-mediated SOX11 induction in the DG (von Wittgenstein et al. 2020) and retinal ganglion cells (RGCs) (Chang et al. 2021). While some of the genes were also down-regulated in the dentate neuroepithelium at E13.5 in the embryos lacking SOX11, specifically in the telencephalon (Abulaiti et al. 2022), the difference was not significant considering these genes as a group. Further supporting the 33 genes being tightly associated with Sox11 activity, Enrichr (Chen et al. 2013) found that they were enriched with genes co-expressed with human SOX11 in the ARCH4 database (adjusted P=2×10^-5^, odds-ratio 16.84). We hereafter refer to these genes as “Sox11 targets”.

**Figure 7.**
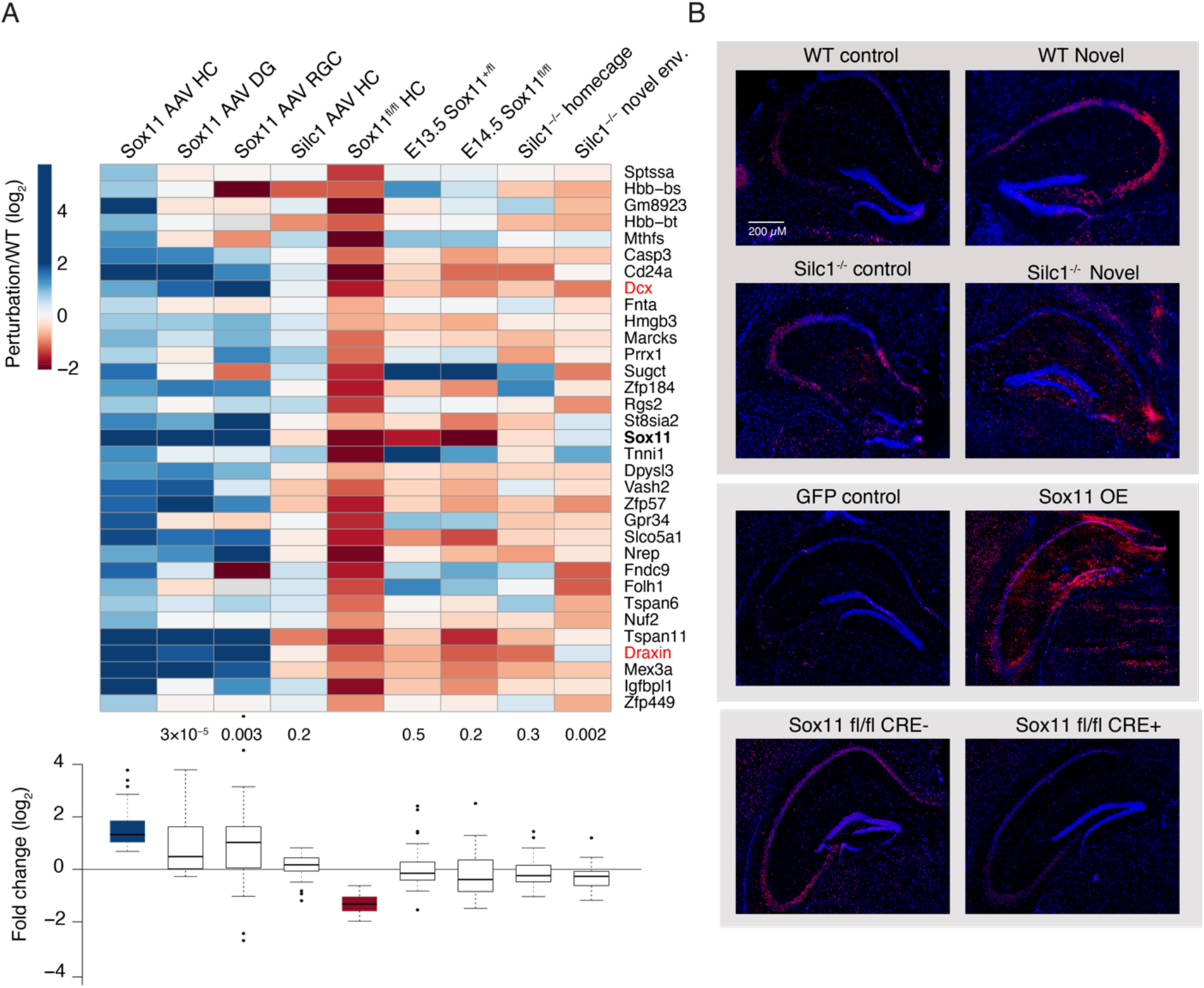
Characterization of genes regulated by *Sox11* and *Silc1* by RNA-seq. **(A)** Top: heatmap of changes in gene expression relative to the respective controls of the 33 genes that were up-regulated following AAV-mediated Sox11 OE in the hippocampus and down-regulated following AAV-mediated introduction of Cre into the hippocampus of *Sox11*^fl/fl^ mice (P<0.05, fold change of at least 50%). Bottom: boxplots of the same changes, with p-values computed using a two-sided one-sample t-test. **(B)** Immunostaining with anti-Draxin (red) and DAPI (blue) in the hippocampi of WT and *Silc1^-/-^* mice, in HC and NE conditions, Sox11 OE and Cre- or Control-injected *Sox11*^fl/fl^ mice. Imaging using 20X objective (Scale bar 200 μm).

Consistently with its inability to lead to changes in SOX11 levels, AAV-mediated induction of *Silc1* did not significantly affect the expression of Sox11 targets (**Fig. 7A**). In contrast, these genes were significantly reduced in the hippocampus of *Silc1^-/-^* mice following NE relative to WT mice. The seven negatively regulated targets were supported in the other datasets to a lesser extent and did not change significantly in the *Silc1^-/-^* hippocampus (**Fig. S8A**).

Another 1,047 genes were down-regulated with the same criteria (25% reduction and P<0.05, **Table S1**) in the E13.5 *Sox11*-null dentate neuroepithelium but did not meet our bi-directional change criteria in the adult hippocampus (“embryonic Sox11 targets”, **Fig. S8B**). Notably, a major fraction of the embryonic Sox11 targets were substantially induced upon Sox11 OE in the DG and RGC but to a much lesser extent in the other section of the hippocampus. While some of the 1,047 genes were down-regulated upon Cre injection, the fold changes were overall modest. In addition, in the *Silc1^-/-^* hippocampus, although the overall reduction was significant, the median fold-change was not substantial, i.e., close to zero. Therefore, we conclude that whereas *Sox11* is required for the proper expression of many genes in the embryo and regulates their expression in immature neurons, only a small subset of genes remains sensitive to SOX11 levels in the mature neurons in the hippocampus. We suggest that in these cells, the expression levels of SOX11 are much lower than in the embryo or in the DG SGZ, therefore even *Sox11* OE via an AAV, which up-regulates *Sox11* by ~30-fold, still results in lower *Sox11* levels than those present in the E13.5 embryo. Alternatively, post-translational modifications of SOX11 in the embryo and the SGZ differ from those in the mature neurons and limit SOX11 activity. Nevertheless, a small yet notable set of specific Sox11 targets, including *Dcx* and *Draxin*, remains sensitive to increases and decreases in Sox11 levels in the mature hippocampus. Crucially, these genes are significantly reduced in the *Silc1^-/-^* hippocampus following exposure to a novel environment (NE condition). We validated changes in DRAXIN protein levels using immunostaining and obtained a similar pattern of results to those of the RNA-seq analysis **(Fig. 7B)**. Collectively, the data suggest that in the adult hippocampus, *Sox11* drives a specific gene expression program containing a subset of the *Sox11* targets in the developing brain; the expression of this program is affected by the loss of *Silc1*. It thus appears that a subset of a regulatory program used during development is co-opted during memory formation in the hippocampus.

**Figure S7.**
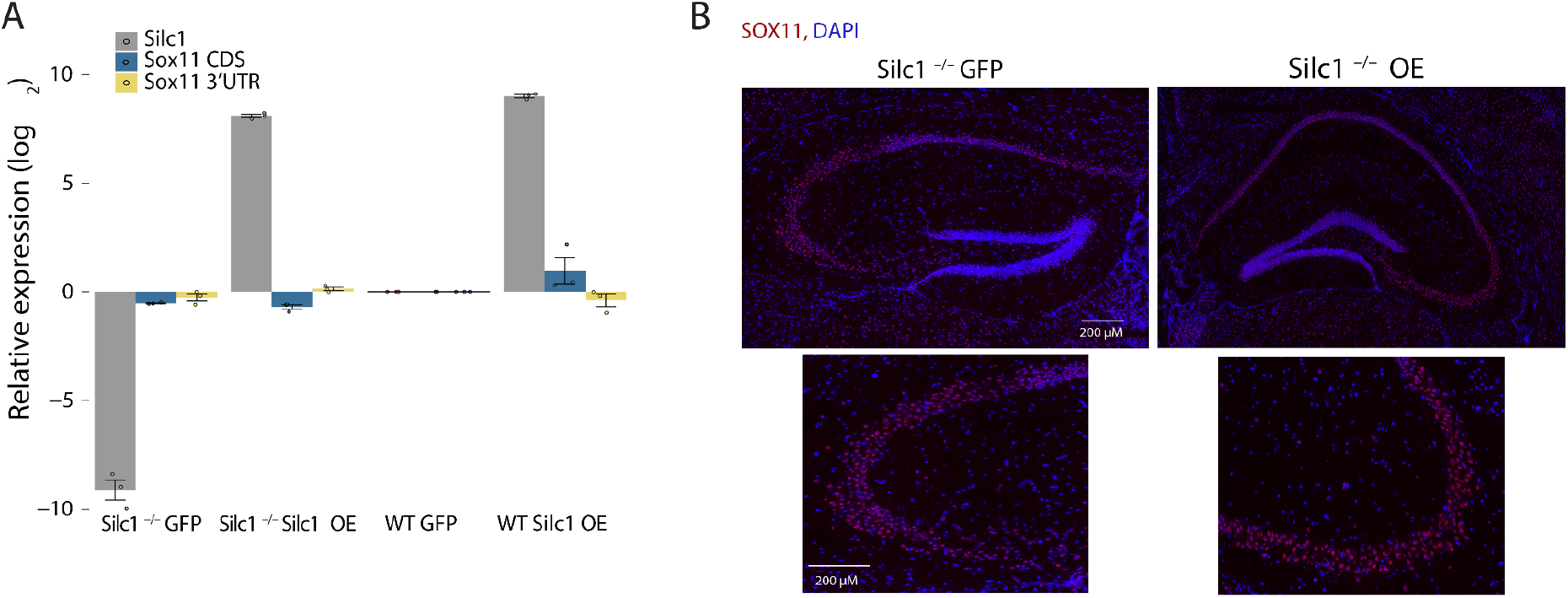
*Silc1* over-expression in *Silc1^-/-^* mice. **(A)** qRT-PCR quantifications of *Silc1* and *Sox11* after injection of AAV9-Silc1 or AAV9-GFP into the CA3 region of WT and *Silc1^-/-^* mice. Levels were normalized to WT mice injected with AAV9-GFP, and β actin was used as an internal control. **(B)** Immunostaining with anti-SOX11 (red) and DAPI (blue) in hippocampi of *Silc1^-/-^* mice injected with Silc1 OE AAV. Imaging using 20X objective (Scale bar 200 μm).

**Figure S8.**
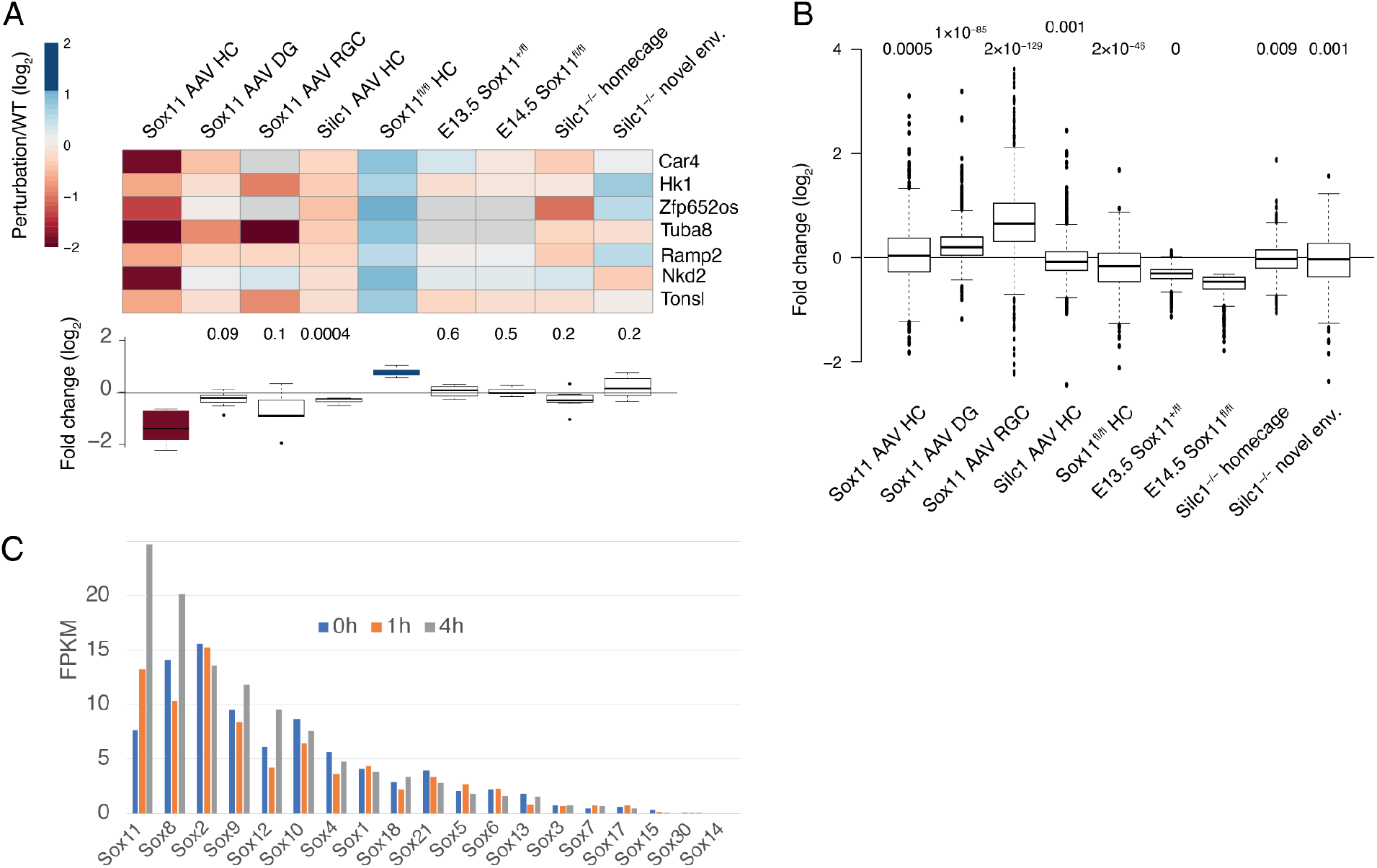
Gene expression in the hippocampus. **(A)** Genes negatively regulated by Sox11 in the hippocampus. As in **Fig. 7**, for the seven genes negatively regulated by Sox11. **(B)** Distribution of the changes in gene expression for the comparisons shown in Fig. 7 for the genes that are significantly reduced in the E13.5 *Sox11^fl/fl^* Cre+ dentate neuroepithelium. P-values for each group obtained using Wilcoxon rank-sum test are shown above each boxplot. **(C)** As in Fig. 1C, for the different genes in the Sox family of TFs. The RNA-seq data are from the mouse dentate gyrus and the indicated time after ECS.

### Loss of *Silc1* results in the reduction of chromatin binding at Sox family binding sites

We presumed that the changes in gene expression are driven by changes in the chromatin and used ATAC-seq (Buenrostro et al. 2013) to profile accessible chromatin in the hippocampus of WT and *Silc1^-/-^* mice placed in a novel environment (six biological replicates). We first quantified accessibility at 31,764 peaks by jointly analyzing the full dataset by MACS2 (Y. Zhang et al. 2008) (**Fig. 8A** and **Table S2**). No peak was differentially accessible between WT and *Silc1^-/-^* mice when accounting at FDR<0.05. The peaks in the two gene deserts flanking Sox11 did not appear to change their accessibility, with the notable exception of an AP-1 bound peak (the central peak in the zoomed-in panel in the figure). To evaluate whether there were changes in transcription factor binding within the accessible regions, we analyzed TF footprinting with TOBIAS (Bentsen et al. 2020). This analysis (**Table S3**) implicated numerous TFs as differentially binding accessible genome regions, notably indicating reduced binding to TF binding sites of the Sox family (**Fig. 8B**), consistent with reduced protein expression of Sox11, which is the main Sox factor expressed in the hippocampus (**Fig. S8C**).

**Fig. 8.**
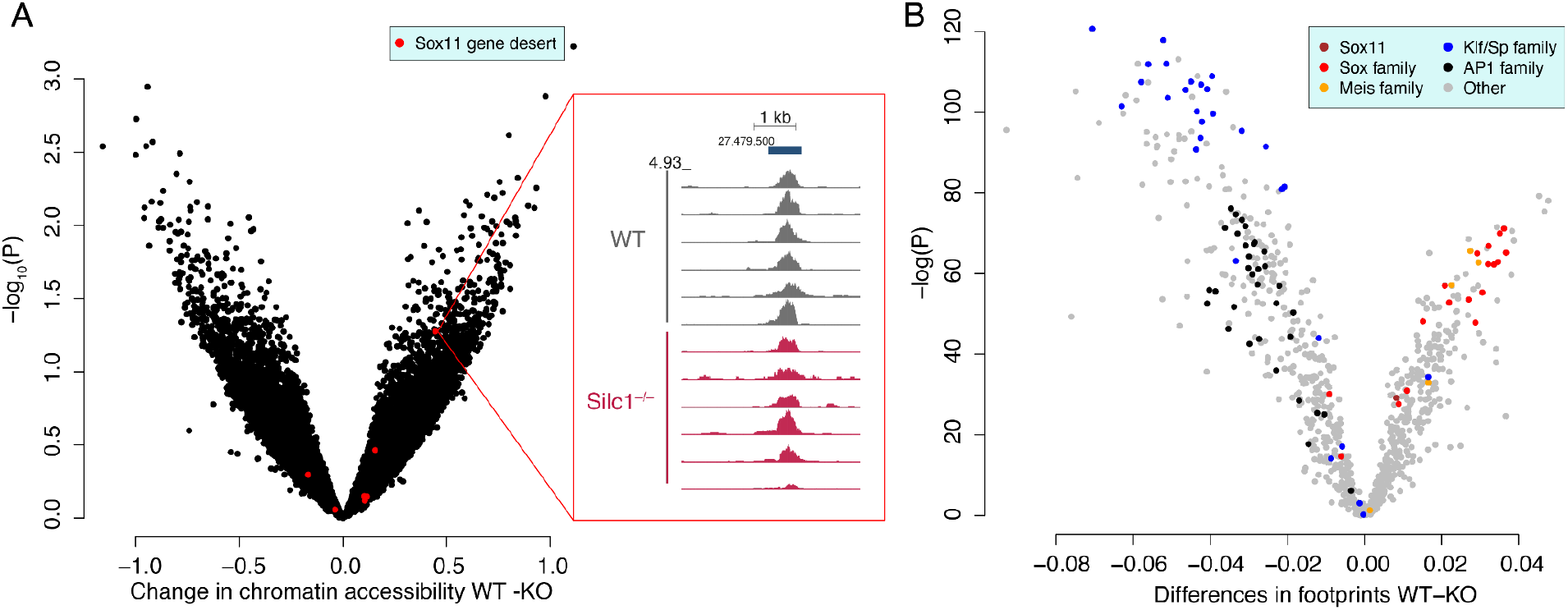
Differences in chromatin accessibility in *Silc1^-/-^* hippocampus. **(A)** Volcano plot of the difference in the read coverage (x-axis) and statistical significance (y-axis) at the 31,764 peaks called by MACS2 using the entire dataset. Peaks in the ~2Mb gene desert flanking Sox11 are in red. The inset shows ATAC-seq read coverage in the most differential Sox11-proximal peak, which is one of the peaks highlighted in **Fig. S1. (B)** Changes in TF footprints for each TF family present in the JASPAR database. Motifs assigned to the indicated families are marked in one of five colors.

### *Silc1* expression is reduced in the aging hippocampus and cortex with further reduction in a model of Alzheimer’s disease

Since *Silc1* is required for timely learning, which is impaired in aging and neurodegenerative diseases, we have further evaluated the effects of aging (4 time points: 4,8,12 and 18 months) and of a hereditary form of age-associated cognitive decline, Alzheimer’s disease (AD) a model. We used data of gene expression collected from the hippocampus and cortex from multiple mice from the C57BL/6 background used in our study (Forner et al. 2021) (**Fig. 9**). These analyses indicated that *Silc1* expression was reduced in both brain regions, with further reduction in the AD model, which expresses five familial AD mutations under the control of a Thy1 mini-gene. In the control mice, *Silc1* expression was strongly negatively correlated with age in both brain regions (hippocampus: R=–0.58, P=6.42×10^-6^ cortex: R=–0.47, P=5.08×10^-4^). At most ages, a further reduction was found in the AD mice relative to controls. These results show that *Silc1* continues to be broadly and abundantly expressed in the forebrain throughout life and that its loss might be associated with aging- and AD-related cognitive decline.

**Figure 9.**
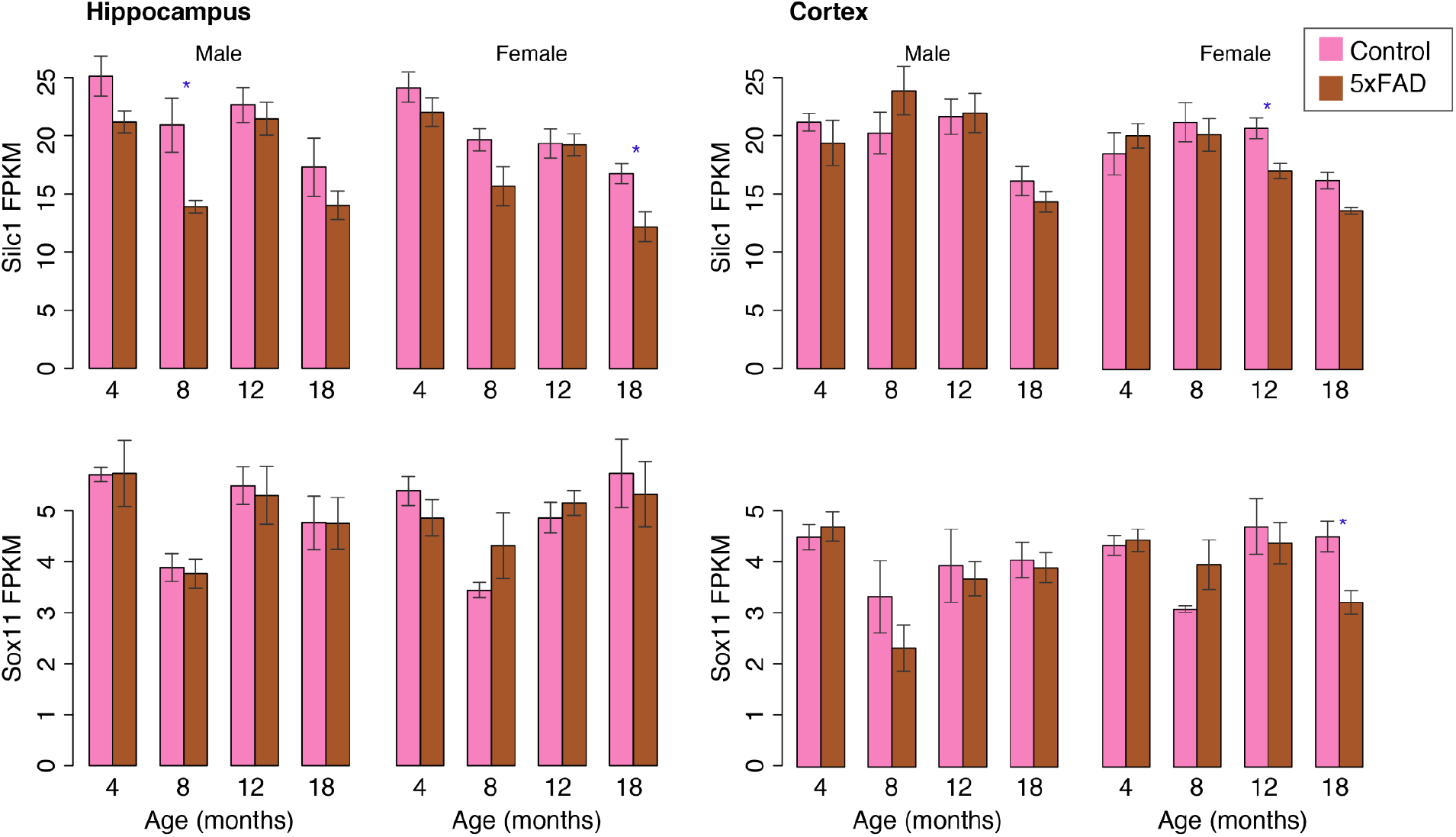
Expression of *Silc1* (top) and *Sox11* (bottom) in the indicated brain region extracted from control WT mice and from 5xFAD mice at the indicated age, data from (Forner et al. 2021). Blue asterisks denote P<0.05 for the comparison between Control and 5xFAD mice, two-sided Wilcoxon rank-sum test.

## Discussion

The mechanism by which *Silc1* facilitates the activation of *Sox11* in the CNS and the PNS remains largely unknown. In both systems, accessibility of the *Sox11* promoter does not appear to change during *Sox11* activation – to the extent measurable by genome-wide ATAC-seq, the promoter appears highly accessible in all neuronal tissues. The paucity of cell lines in which *Silc1* is endogenously expressed hinders some experimental approaches for its perturbation. *Silc1* is not expressed in mouse cell lines, the mouse embryo, or in commonly used primary hippocampal cultures obtained from embryonic or early postnatal stages. We have managed to successfully obtain and characterize, by qPCR and RNA-seq, primary cultures of adult hippocampal neurons but found that those cells lost all expression of *Silc1* or *Sox11* (**Fig. S5A**). While Neuro2a cells express some *Sox11* and can be induced by CRISPR activation to express *Silc1* (Perry et al. 2018), these levels of induction are insufficient to drive levels of *Sox11* expression comparable to those in the brain. Therefore, we are presently limited to experimental manipulations that can be performed in the adult brain, such as the introduction of GapmeRs and AAVs as performed here.

Notably, while *Silc1* has some effect on the basal levels of *Sox11* expression in the nervous system, which is low and lower than those of *Silc1*, it appears to be particularly important for the timely induction of *Sox11* expression upon different physiological cues. As such, it is reminiscent of other *cis*-acting RNAs (Gil and Ulitsky 2020), the modes of action of which are largely unknown. A common denominator of these RNAs is a relatively high expression in unstimulated conditions and efficient splicing, which has been linked with more efficient enhancer activity in broad regions flanking the spliced lncRNAs (Gil and Ulitsky 2018; Jennifer Y. Tan et al. 2020; Jennifer Yihong Tan and Marques 2022). It is, therefore, possible that transcription of *Silc1* and its splicing enables the proper positioning of the broad *Sox11* locus in a nuclear and chromatin environment that facilitates a stronger response to stimulus, such as that which occurs when mice are exposed to NE. Indeed, we found a region that corresponds to an AP-1–regulated enhancer as differentially accessible in the *Silc1^-/-^* hippocampus. However, these changes were variable between mice and do not reach statistical significance (**Fig. 8**). Development and availability of methods that will allow measurement of chromatin accessibility *in situ* can potentially shed further light on whether this enhancer becomes specifically less accessible in cells that normally activate *Sox11* expression following exposure to NE.

Our analysis of the available atlases of gene expression has shown partially overlapping domains of expression of the two genes. In principle, single-cell RNA-seq (scRNA-seq) data can provide some additional insight, but we found that the limited expression levels (FPKM of 1–10) of both genes limit the number of cells in which both of them can be detected. For example, even though we see a rather high expression and potent induction of *Silc1* expression with RNAscope (**Fig. 1B** and **Fig. 3A**), it is difficult to observe these trends in publically available data on scRNA-seq expression in the hippocampus following NE exposure (Lacar et al. 2016), as individual cells sequenced in that experiment had very few reads originating from *Silc1* locus. Similarly, whereas single-cell datasets from the dentate gyrus supported expression of *Sox11* in immature neurons (http://linnarssonlab.org/dentate/), there were very few reads from other neurons, which evidently express *Sox11* based on RNAscope imaging.

We focused here on the interplay between *Sox11* and *Silc1* in the hippocampus and on their roles in spatial memory acquisition, but notably, the two genes, and in particular, *Silc1*, are broadly expressed in the adult brain, suggesting that in certain conditions, *Silc1* can be important outside of the hippocampus. We focused on the hippocampus as electroconvulsive stimulation of the whole brain most prominently activates *Sox11* in the DG (Sun et al. 2005), and we are not aware of other physiological conditions where *Sox11* is transcriptionally activated in the adult brain. It is possible that other conditions require activation of the “immature neuron” transcriptional program, including *Dcx*, *Draxin*, and the other *Sox11* targets we describe here.

*Sox11* has been studied in many systems, including the retina, but within the forebrain, it has been studied almost exclusively in the context of the developing brain or the immature neurons in the SGZ. The view emerging from multiple studies is that in these systems, *Sox11* regulates targets shared in part with those of *Sox4* and supports further maturation of neurons that recently exited the cell cycle. Surprisingly, we find that in NE conditions, *Sox11* is prominently induced in mature parts of the hippocampus. Among the genes differentially expressed by loss of *Silc1*, which presumably affects predominantly mature neurons, as does not appear to be expressed in immature ones, we find a small but prominent set of *Sox11* targets that are regulated by *Sox11* also in the context of immature neurons, and normally also predominantly expressed in neuronal progenitors and immature neurons in the SGZ, such as *Dcx*, *Draxin* (S. Zhang et al. 2010), and *Prrx1* (Shimozaki, Clemenson, and Gage 2013).

These appear to be a subset of the genes that depend on *Sox11* in the developing brain, though many of those targets might be indirect. The design principle underlying this apparent re-use of a master regulator of neuronal maturation in the context of plasticity of mature neurons will be an interesting subject for future studies. Interestingly, the loss of some of the *Sox11* targets prominently regulated is related to abnormal mossy fiber formation (*St8sia2* (Angata et al. 2004)). Notably, among the *Sox11* targets, loss of *Dcx* as well as P311, encoded by the *Nrep* gene leads to defects in spatial learning (Taylor et al. 2008; Corbo et al. 2002).

Combined with our previous observations in the PNS, this study solidifies the notion that *Sox11* has at least two main regulatory regimes. The first is active in immature neurons, in the embryo and in the SGZ. These are cells that exited the cell cycle but undergo neuronal growth, which is apparently dependent on high SOX11 levels. Presumably, a variety of enhancer elements are supporting this high *Sox11* expression, as neither *Silc1* nor other lncRNAs in the large gene deserts flanking *Sox11* appear to be expressed in these cells. The second regime is activated in cells that cease their growth and possibly coincides with the expression of *Thy1*, which is the gene most tightly correlated with *Silc1* in the mouse NeuroSeq expression atlas. In basal conditions, this program supports minimal expression of *Sox11*, which is likely not required for the steady-state activity of these neurons. Upon activation, likely via the activity of the AP-1 TFs, *Sox11* is induced quite potently in this regime. This activation can result from injury signaling in the PNS, or from exposure to NE in the CA subfields in the hippocampus. A timely and potent induction of *Sox11* in these conditions depends on *Silc1* transcription or RNA product, as it is sensitive to perturbation by antisense oligonucleotides.

## Materials and Methods

### Animals

The study was conducted following the guidelines of the Weizmann Institutional Animal Care and Use Committee (IACUC). C57black6 Ola HSD male mice were purchased from Harlan Laboratories (Rehovot, Israel). All other mouse strains were bred and maintained at the Veterinary Resources Department of the Weizmann Institute.

### DRG cultures

Adult mouse DRGs were dissociated for neuron cultures with 100 U of papain followed by 1 mg/ml collagenase-II and 1.2 mg/ml dispase. The ganglia were then triturated in HBSS, 10 mM glucose, and 5 mM HEPES (pH 7.35). Neurons were recovered through percoll, plated on laminin, and grown in F12 medium for 48 hours (Hanz et al., 2003; Perlson et al., 2005; Rishal et al., 2010). Adult male mice DRG cultures were transfected with GapmeRs using DharmaFect 4 (Dharmacon). 72 hr after transfection total RNA was extracted to ensure knockdown.

### Generation of *Silc1* polyA mice

Mice carrying a *Silc1*^polyA^ allele were generated using the CRISPR/Cas9 system for insertion of transcription terminator by standard procedures at the Weizmann transgenic core facility using a single guide RNA (sgRNA) targeted to exon 1 (chr12:27160402, mm10 assembly). gRNA sequence was designed using CHOPCHOP (Labun et al. 2016) and ordered from IDT (gRNA sequence: GTGCTTGGCACTGCTTGGCA). For homologous recombination, an ssODN (200 nt) containing two homology arms (50 nt each), a short poly(A) site (49 nt), and two MAZ sites (Ballarino et al. 2018) was synthetized by IDT (**Table S4**). The poly(A)/MAZ insertion was detected by PCR amplifications. Sequences of primers used for genotyping appear in **Table S4**. Lines were bred and maintained on C57BL/6 background at the Veterinary Resources facility of the Weizmann Institute. All the experiments were done on 6–8 weeks old mice from F3 generation.

### Generation of *Silc1* conditional mice

Mice carrying the *Silc1^fl^* conditional alleles were generated using the CLICK system (Miyasaka et al. 2018) using a long single-stranded DNA (lssDNA), by standard procedures at the Weizmann transgenic core facility. Two single guide RNAs (sgRNAs) targeted to sites before the *Silc1* promoter (chr12:27160159, mm10 assembly) and after exon 1 (chr12:27161872, mm10 assembly) were designed using CHOPCHOP (Labun et al. 2016) and ordered from IDT (Perry et al. 2018)). 12 ug of the lssDNA were also ordered from IDT. *Silc1^fl/fl^* mice were identified by genotyping and sequencing using primers flanking the loxP insertion sites. Sequences of primers used for genotyping appear in **Table S4.** Lines were bred and maintained on C57BL/6 background at the Veterinary Resources facility of the Weizmann Institute. All the experiments were done on 6–8 weeks old mice.

### Morris Water Maze

The water maze (Nunez 2008) consisted of a circular tank (120 cm diameter) filled with 24°C water clouded with milk powder with a removable escape platform centered in one of the four maze quadrants. In the testing room, only distal visual-spatial cues for locating the hidden platform were available. *Acquisition phase* - The mice were subjected to 4 trials per day with an inter-trial interval of 2 minutes, for 8 consecutive days. In each trial, the mice were required to find a platform located in one of the four quadrants submerged 0.5 cm below the water surface within 90 sec. The escape latency in each trial was recorded. Each mouse was allowed to remain on the platform for 15 s and was then removed from the maze. If the mouse did not find the platform in the allocated time, it was manually placed on the platform for 15 s. *Probe test* - Memory was assessed 24 hours after the last trial. The escape platform was removed, and mice were allowed to search for it for 1 minute. The time spent, the swimming distance in the different quadrants of the pool, and the time spent (percentage), were monitored using an automated tracking system (Ethovision XT, Noldus, the Netherlands).

### Barnes circular maze

The test was performed as previously described (Pitts 2018). The apparatus used was an elevated circular platform (0.91 m in diameter) with 20 holes (5 cm diameter) around the perimeter of the platform, one of which was connected to a dark escape recessed chamber (target box). The maze was positioned in a room with large, simple visual cues attached to the surrounding walls. Mice were habituated to the training room before each training day for 1 hour in their cage. The acquisition consisted of four daily trials for 4 days, separated by a 15 min intertrial interval. Each mouse was positioned in the center of the maze in an opaque cylinder for 1 min, which was gently lifted and removed to start the session. The mice were allowed for 3 min to find the target box. At the end of the 3 min, if the mouse failed to find the recessed escape box, it was gently guided to the chamber and allowed to stay in the target platform for 1 min. The location of the escape box was kept constant with respect to the visual cues. An animal was considered to enter the escape chamber when the animal’s entire body was inside the chamber and no longer visible on the platform. Retention was tested 24 h after the last training session (day 5); in the probe trial, the target hole was closed, and the latency to reach the virtually target hole was measured. The same parameters were collected during the acquisition and retention phases using an automated tracking system (Ethovision XT, Noldus, the Netherlands).

### Fear conditioning

The test was performed as previously described (Sharma et al. 2018). A computer-controlled fear-conditioning system (Ethovision XT, Noldus, the Netherlands) monitors the procedure while measuring inactivity (freezing) behavior. *1. Conditioning:* conditioning takes place on day 2 in one 5-min training session. Mice were placed in the chamber to explore the context for 2 min. Then we applied a conditioned stimulus (CS) for 30 s, 3,000 Hz, pulsed 10 Hz, 80 dB, and a foot shock for an unconditional stimulus (US): 0.7 mA, 2 s, constant current. The CS–US pairing was repeated twice with a fixed inter-trial interval (ITI) of 60 sec. The US is delivered through the metal grid floor. Mice were removed from this chamber 1 min after the last CS-US pairing and put back in their home cage. A constant auditory background noise (white noise, 62 dB) was presented throughout the experiment. *2. Testing:* Context-dependent memory was tested 24 h after the conditioning by re-exposure to the conditioning box for 5 min without any CS. The Cue dependent memory was tested 1 h after the Context test by exposure to the CS in different environmental conditions (black Plexiglas box, black floor instead of metal grid, no illumination, no background noise, cleaning solution: acetic acid 10% instead of alcohol 10%).

### Western blot and immunofluorescence

Brain sections were fixed with 4% paraformaldehyde for 3 hr followed by overnight in 30% sucrose with overhead rotation. Tissue was frozen in Tissue-Tek O.C.T compound (Sakura 4583) blocks and sectioned using a Leica cryostat (CM3050) at 10 micrometers thickness. Blocking and permeabilization were done with 5% donkey serum, 2% BSA, and 0.1% Triton X-100 in PBS. Primary antibodies were diluted in a permeabilization buffer. Antibodies used: Sox11 antibody, anti-rabbit (ABN105) from Millipore, NeuN antibody, anti-mouse (MAB377) from Millipore and Draxin antibody, anti-rabbit (ab117452) from Abcam. Secondary antibodies: Donkey anti-Mouse Alexa 594 (Molecular Probes A21203) and Goat anti Rabbit Alexa 647 (Abcam ab150079). Nuclei were stained using DAPI (Thermo Fisher Scientific). Imaging was done using a Leica DM4000 B microscope with Leica DFC365 FX CCD camera and Leica application suite (LAS) X software. Western blots were carried out as previously described (Hanz et al., 2003; Perlson et al., 2005). For Westerns, the samples were resolved on 10% SDS PAGE, transferred to nitrocellulose, and incubated with primary antibodies overnight. Antibodies used: Sox11 antibody, anti-rabbit (ABN105) from Millipore and beta-tubulin antibody, antimouse (T4026) from sigma. AzureSpectra fluorescent 700 anti-mouse and 800 anti-rabbit (Azure biosystem) were used as the secondary antibodies for fluorescent quantification of Western blots. Blots were imaged on an Azure Imager system.

### Quantification of immunofluorescence

The immunofluorescence staining was quantified using Fiji (ImageJ) analysis software.

### RNAscope FISH

Brains were immediately frozen on dry ice in tissue-freezing medium. Brains were sliced on a cryostat (Leica CM 1950) into 8-μm sections, adhered to SuperFrost Plus slides (VWR), and immediately stored at −80 °C until use. Samples were processed according to the ACD RNAscope Fluorescent Multiplex Assay manual using Silc1 probes-RNAscope 2.5vs probe “Mm GM9866” Cat No. 536709, Sox11 CDS probe Cat No. 440811, Sox11 3’UTR probe Cat No. 805071 and Fos probe Cat No. 316921. Imaging was performed on a Nikon-Ti-E inverted fluorescence microscope with a 100× oil-immersion objective and a Photometrics Pixis 1024 CCD camera using MetaMorph software as previously described (Bahar Halpern and Itzkovitz 2016).

### RNAscope Quantification

RNAscope analysis was done using IMARIS (v7.7.2) software.

### RNA extraction and sequencing

Total RNA was extracted from the hippocampus using the TRIREAGENT (MRC) according to the manufacturer’s protocol. Strand-specific mRNA-seq libraries were prepared from 1 ug total RNA using the SENSE-mRNA-Seq-V2 (Lexogen), according to the manufacturers’ protocol and sequenced on a NextSeq 500 machine or Novaseq 6000 machine to obtain 75 nt and 150 nt single- or paired-end reads. All RNA-seq dataset is deposited in GEO database with the accession GSE216643.

### Quantitative reverse-transcription PCR (qRT-PCR)

Reverse transcription was done using qScript Flex cDNA synthesis kit (Quanta Biosciences), using random primers. Quantitative PCR was performed in a ViiA 7 Real-Time PCR System (Thermo) in a 10 μl reaction mixture containing 0.1 μM forward and reverse primers, fast SYBR master mix (Applied Biosystems), and template cDNA. A reaction containing DDW instead of cDNA was used as a no-template control and was amplified for each primer pair. Only samples free of DNA contamination were further analyzed. The gene-specific primer pairs used for mRNA expression level analysis are listed in **Table S4**.

### ATAC-seq

ATAC-seq was performed as described (Buenrostro et al. 2013) with minor adjustments for brain tissue. Briefly, hippocampus tissue was extracted in 500 μl Nuclear extraction buffer (10mM Tris, 10mM NaCl, 3mM MgCl2, 0.1% Igepal, 0.1% Tween, protease inhibitor cocktail) for 5 minutes on ice, then a 21g needle on a 1 mL syringe was used to shear the tissue through the needle 5 times. NeuN-positive nuclei were separated by fluorescence-activated cell sorting. Libraries were sequenced with 50 bp paired-end mode on NovaSeq6000.

### Microinfusion of antisense LNA GapmeR

8 weeks old C57BL/6J male mice (Envigo, Israel) were anesthetized with isoflurane and placed in a stereotactic frame (n =3 per group). The skull was exposed to antiseptic conditions and a small craniotomy was made with a thin drill over the hippocampus. Antisense LNA GapmeRs (custom designed, 3’-FAM-labeled, Qiagen, **Table S5**) were bilaterally microinfused using a 2 μl calibrated micropipette (Hamilton syringes ga 25/70mm/pst3), which was pulled to create a long narrow shank. 1 μl was infused slowly by pressure infusion into the CA3 region (from Bregma +3.1 mm anteroposterior, ±2.8 mm mediolateral and +3.2 mm dorsoventral axis), the micropipette was kept in place for 30 s to ensure adequate diffusion. The wound was sutured with sterile nylon material.

### AAV9 vectors cloning and virus generation

AAV9 expression of GFP-Cre from CamKII promoter: Addgene number-105551-AAV9 (plasmid: pENN.AAV.CamkII.HI.GFP-Cre.WPRE.SV40) and AAV9 control virus - 105541-AAV9 (plasmid: pENN.AAV.CamKII0.4.eGFP.WPRE.rBG).

AAV9 plasmids were used for Silc1 and Sox11 overexpression. We cloned Silc1 from The Silc1 pcDNA3.1(+) vector (Perry et al. 2018) and Sox11 from pLenti-CMV-GFP-Sox11 vector (Addgene: #120387) into pENN.AAV.CamKII0.4.eGFP.WPRE.rBG plasmid downstream of the GFP sequence. Recombinant AAV9 plasmids were produced by transfecting HEK293T cells using the AAVpro helper-free systems. AAV9 viral preparations were purified using the AAVpro® Purification Kit (Takara Bio. Inc., Cat#6666).

### Microinfusion of AAV9 viruses

8 weeks-old C57BL/6J male mice (Envigo, Israel, n =3 per group) received bilateral stereotaxic injections of AAV9 Silc1, AAV9 Sox11 or AAV9 GFP control into the hippocampus CA3 regions (titer of 10^12^ vg/ml, 0.2 μl Min). The virus was delivered using a 2 μl Hamilton syringe connected to a motorized nano-injector. To allow diffusion of the solution into the brain tissue, the needle was left in place for 4 min after the injection (from Bregma +3.1 mm anteroposterior, ±2.8 mm mediolateral and +3.2 mm dorsoventral axis). The wound was sutured with sterile nylon material. The mice recovered from the surgery for a period of 2-3 weeks before the hippocampus was extracted for RNA extraction or for sections. The slides were screened for GFP signal at the injection site and mice that did not show fluorescent labeling at the aimed injection location were excluded from the data. AAV9 Cre-GFP and AAV9 GFP were injected into Sox^fl/fl^ mice and Silc1^fl/fl^ Mice using the same parameters.

### RNA-seq data analysis

RNA-seq reads were mapped to the mouse genome (mm10 assembly) using STAR (Dobin et al. 2013) to generate read coverage tracks visualized using the UCSC genome browser. RSEM (Li and Dewey 2011) with RefSeq annotations was used to call differential expression between samples with data collected in this study and data from public datasets on OEof Sox11 in the hippocampus: SOX11 induction in the DG (von Wittgenstein et al. 2020) – SRP229390; SOX11 induction and in RGCs (Chang et al. 2021) – SRP290800; dentate neuroepithelium at E13.5 in the embryos lacking SOX11 (Abulaiti et al. 2022) – SRP285830; aging hippocampus and cortex – SRP309056. Differential expression between conditions was called using DESeq2 with default parameters (Love, Anders, and Huber 2014).

### ATAC-seq data analysis

ATAC-seq reads were mapped to the mouse genome using Bowtie2 (Langmead and Salzberg 2012) and peaks were called using all the samples together using MACS2 (Y. Zhang et al. 2008). The peaks were then adjusted to a fixed width of 140 nt around the peak summit and differential read coverage between the six WT and the six Silc1^-/-^ samples was called using HOMER (Heinz et al. 2010) with the default DESeq2 parameters. The data were also processed using TOBIAS (Bentsen et al. 2020) with default parameters to compute differential TF footprints.

## Data availability statement

All RNA-seq and ATAC-seq data are deposited in GEO database with the accession GSE216643 (reviewer token krylqcsmpdwvvod).

## Supplemental Item Titles

**Table S1. Changes in gene expression computed by DESeq2 in the datasets produced and re-analyzed in this study.**

**Table S2. Accessibility and changes in accessibility in ATAC-seq data from the hippocampus of mice placed in a novel environment.**

**Table S3. Results of the TOBIAS analysis of the ATAC-seq data.**

**Table S4. Primers**

**Table S5. GapmeRs sequences**

## Acknowledgments

We thank members of the Ulitsky laboratory for helpful discussions and comments on the manuscript; Liat Fellus Alyagor and Dana Hirsch for help with RNAscope experiments and analysis; Dieter Chichung Lie, Yaniv Ziv, Ofer Yizhar, and Ivo Spiegel for helpful discussions. Sox11 loxP mice were a kind gift from Prof. Veronique Lefebvre (Leonard and Madlyn Abramson Pediatric Research Center, Philadelphia, USA).

## Author Contributions

RBP and IU conceived the study. RBP carried out all the experiments with assistance from M. Tsoory with behavioral experiments and M. Tamasov that performed the stereotaxic injections. IU analyzed the high-throughput sequencing data and RBP analyzed all other data. RBP and IU wrote the manuscript.

## Declaration of Interests

The authors declare no competing interests.

## Notes

### Competing Interest Statement

The authors have declared no competing interest.

